# CRISPR-Cas systems are widespread accessory elements across bacterial and archaeal plasmids

**DOI:** 10.1101/2021.06.04.447074

**Authors:** Rafael Pinilla-Redondo, Jakob Russel, David Mayo-Muñoz, Shiraz A. Shah, Roger A. Garrett, Joseph Nesme, Jonas S. Madsen, Peter C. Fineran, Søren J. Sørensen

## Abstract

Many prokaryotes encode CRISPR-Cas systems as immune protection against mobile genetic elements (MGEs), yet, a number of MGEs also harbor CRISPR-Cas components. With a few exceptions, CRISPR-Cas loci encoded on MGEs are uncharted and a comprehensive analysis of their distribution, prevalence, diversity, and function is lacking. Here, we systematically investigated CRISPR-Cas loci across the largest curated collection of natural bacterial and archaeal plasmids. CRISPR-Cas loci are widely but heterogeneously distributed across plasmids and, in comparison to host chromosomes, their mean prevalence per Mbp is higher and their distribution is markedly distinct. Furthermore, the spacer content of plasmid CRISPRs exhibits a strong targeting bias towards other plasmids, while chromosomal arrays are enriched with virus-targeting spacers. These contrasting targeting preferences dominate across the diversity of CRISPR-Cas subtypes and host taxa, highlighting the genetic independence of plasmids and suggesting a major role of CRISPR-Cas for mediating plasmid-plasmid conflicts. Altogether, CRISPR-Cas are frequent accessory components of many plasmids, which is an overlooked phenomenon that possibly facilitates their dissemination across microbiomes.

## INTRODUCTION

Clustered regularly interspaced short palindromic repeats (CRISPR) and their associated (*cas*) genes encode adaptive immune systems that provide prokaryotes with sequence-specific protection against viruses, plasmids, and other mobile genetic elements (MGEs) (Barrangou and Marraffini, 2014). These systems consist of two main components: 1) a CRISPR array, which is a DNA memory bank composed of sequences derived from previous infections by MGEs, and 2) *cas* genes that encode the protein machinery that is necessary for the three stages of immunity (adaptation, RNA biogenesis, and interference) (Hille *et al*., 2018). Briefly, during adaptation, short sequence fragments from the genomes of invading MGEs are integrated at the CRISPR leader end as new “spacers” flanked directly by repeats in the array. Biogenesis involves expression of the CRISPR array as a long transcript (pre-crRNA) and its subsequent processing into mature CRISPR RNAs (crRNAs), each corresponding to a single spacer. Finally, during interference, the mature crRNAs are coupled with one or multiple Cas proteins in search of a complementary sequence (protospacer), leading to the nuclease-dependent degradation of target nucleic acids.

CRISPR-Cas systems are broadly distributed across the genomes of about 42% of bacteria and 85% of archaea (Makarova *et al*., 2020). Despite the aforementioned commonalities, these systems display remarkable diversity in their mechanisms of action and in the phylogeny of their components. They are divided into two major classes, six types and more than 45 subtypes on the basis of the distinct architectures and the organization of their effector modules (Pinilla-Redondo *et al*., 2019; Makarova *et al*., 2020). Previous work has focused primarily on investigating the canonical adaptive immune functions of CRISPR-Cas systems, their distributions across prokaryotic lineages, and their numerous biotechnological applications (Barrangou and Doudna, 2016; Pickar-Oliver and Gersbach, 2019). Although much less attention has been paid to their presence and function in MGEs, recent research demonstrates that CRISPR-Cas loci are encoded by different types of MGEs (Faure *et al*., 2019). Several viruses, transposons, and plasmids have been shown to carry CRISPR-Cas components that perform different roles, including participating in inter-MGE warfare (Crowley *et al*., 2019; McKitterick *et al*., 2019; Medvedeva *et al*., 2019; Nasko *et al*., 2019; Pinilla-Redondo *et al*., 2019; Al-Shayeb *et al*., 2020), RNA-guided DNA transposition (Peters *et al*., 2017; Klompe *et al*., 2019; Strecker *et al*., 2019), and in anti-defense functions (Seed *et al*., 2013; Faure *et al*., 2019).

Plasmids are extrachromosomal, self-replicating MGEs that are ubiquitous across microbiomes on Earth. They are known to shape the ecology and evolution of microbial communities by, for example, promoting horizontal gene transfer (HGT) between taxa (Norman, Hansen and Sørensen, 2009; Harrison and Brockhurst, 2012). Although the fates of plasmids are linked to those of their microbial hosts, plasmids and host chromosomes are subject to distinct selective constraints and follow different evolutionary trajectories (Lili, Britton and Feil, 2007; MacLean and San Millan, 2015). Despite the beneficial traits that some plasmids provide to their hosts under certain conditions (e.g. antibiotic or heavy metal resistance), they can also impose a physiological burden. Thus, plasmid-host relationships are often dynamic and, depending on the ecological context, extend from parasitic to mutualistic (MacLean and San Millan, 2015). Epitomizing the existence of plasmid-host conflicts, a fraction of chromosomal CRISPR spacers typically match plasmids (Touchon and Rocha, 2010; Shmakov *et al*., 2017), more frequently at specific regions (i.e. the leading strand of conjugative plasmids) (Westra *et al*., 2013). Furthermore, several studies have reported experimental evidence for strong CRISPR-based anti-plasmid immunity (Marraffini and Sontheimer, 2008; Garneau *et al*., 2010; Hatoum-Aslan *et al*., 2014). In turn, many plasmids carry Anti-CRISPR proteins that block host CRISPR-Cas targeting (Mahendra *et al*., 2020; Rafael Pinilla-Redondo *et al*., 2020).

Even though some plasmids have been reported to encode CRISPR-Cas loci (Godde and Bickerton, 2006; Millen *et al*., 2012; Lange *et al*., 2013; Scholz *et al*., 2013; Maier, Dyall-Smith and Marchfelder, 2015; Faure *et al*., 2019; McDonald *et al*., 2019; Özcan *et al*., 2019; Bernheim *et al*., 2020) their incidence, diversity, distribution and function(s) remain largely unstudied. Type IV CRISPR-Cas systems, in particular, are found almost exclusively on plasmids (Faure *et al*., 2019; Faure, Makarova and Koonin, 2019; Makarova *et al*., 2020) and recent work indicates that they participate in plasmid-plasmid competition dynamics (Crowley *et al*., 2019; Pinilla-Redondo *et al*., 2019). Furthermore, a study analyzing CRISPR-Cas systems across a large subset of prokaryotic genomes identified several plasmid-encoded CRISPR-Cas loci, whereas very few were encoded by associated (pro)phages (Bernheim *et al*., 2020). Here, we undertook the first systematic investigation of CRISPR-Cas contents across publicly available bacterial and archaeal plasmid data. We focused on analysing their prevalence, distribution and diversity, and investigated their CRISPR array spacer contents to infer their biological functions.

## RESULTS

### 1. CRISPR-Cas systems are common on plasmids

We scanned the largest curated collection of complete wildtype bacterial (27,939) and archaeal (253) plasmid genomes in search of CRISPR and *cas* loci. To reduce the confounding effect of sequencing biases, we removed identical or highly similar plasmids from further analyses. This resulted in a non-redundant dataset of 17,608 bacterial and 220 archaeal plasmid sequences, spanning 30 phyla and 771 genera. For a total of 13,265 non-redundant plasmids, we were able to collect the corresponding set of host chromosome sequences (n=6,979). Overall, our survey identified a total of 338 complete and 313 putatively incomplete loci (207 orphan CRISPR arrays and 106 orphan *cas*), indicating that ~3% of sequenced plasmids naturally carry one or more CRISPR and/or *cas* loci (Figure 1A, top). This contrasts with the much higher incidence we found on the plasmid-associated host chromosomes, which amounted to 42.3% (42% in bacteria and 63% in archaea). However, since chromosomes are substantially larger than plasmids, we corrected their incidence to genome sequence length (per Mbp) (Madsen *et al*., 2018). Strikingly, we found that CRISPR-Cas components are on average more prevalent across plasmid sequences (Figure 1A, bottom), suggesting a selective advantage for many plasmids to carry these systems.

**Figure 1.**
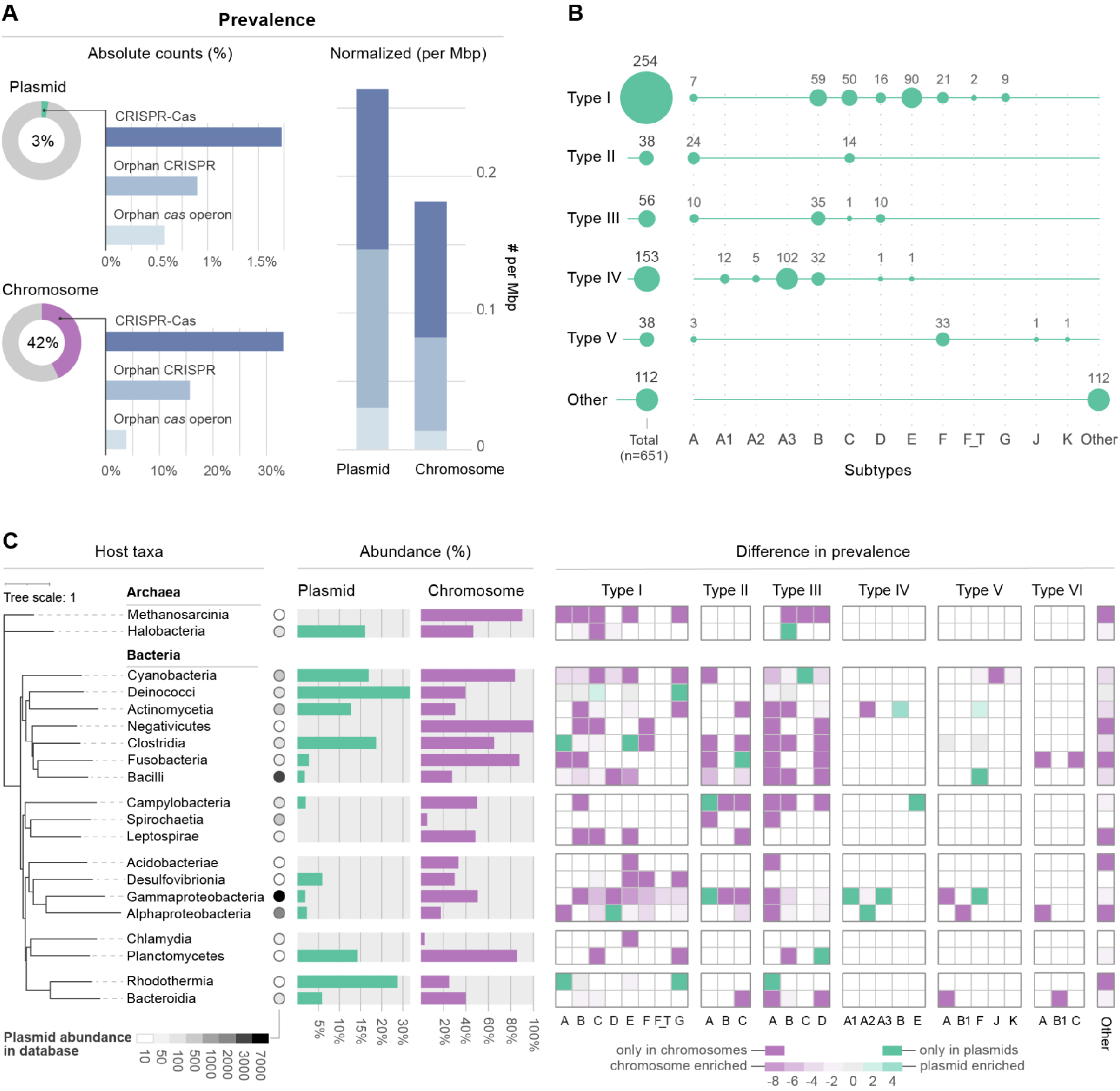
Prevalence, diversity and distribution of CRISPR-Cas loci across plasmid genomes. **(A)** Prevalence of CRISPR-Cas components encoded on non-redundant plasmid genomes and plasmid-associated host chromosomes, expressed as absolute counts per replicon (left) and normalized to counts Mbp (right). **(B)** Distribution and incidence of CRISPR-Cas subtypes across non-redundant plasmids; “other” represents systems that could not be unambiguously assigned (e.g. co-localised/hybrid systems and orphan-untyped components). “I-F_T” refers to the transposon-associated subtype I-F variant and subtype IV-A is subdivided into its known variants (IV-A1 to A3). Total counts per CRISPR-Cas type are summarised on the left. **(C)** Prevalence and distribution of CRISPR-Cas loci in plasmid genomes (green) and in plasmid-host chromosomes (purple) across taxa (at class level). Prevalence is expressed as the percentage of replicons carrying CRISPR-Cas. The lane of circles reflects the abundance of publicly available plasmid genome sequences for each plasmid-host class. Taxa are displayed at the tips of a neighbour-joining tree that is built on the basis of the median cophenetic distance (from the whole-genome GTDB phylogeny) between the different classes and rooted by the archaeal clade. The taxonomic differences in prevalence of the different CRISPR-Cas subtypes are displayed as a heat map with log2 ratios between plasmid and chromosome prevalence. Only taxa for which at least 10 non-redundant plasmids are characterised are displayed.

Whereas most detected loci represent complete CRISPR-Cas systems, solitary (orphan) CRISPR arrays and *cas* operons were also commonly identified. These putatively incomplete systems are more frequent on plasmids than chromosomes (Figure 1A). Intriguingly, the average lengths of orphan arrays are significantly smaller than *cas*-associated CRISPRs (on average 39% shorter, P<2e-16, negative-binomial generalized linear model; Supplementary Figure S1A), which may reflect the importance of neighboring adaptation modules (*cas1-2*) for array expansion and maintenance. Furthermore, we found a lower association of plasmid-encoded systems with adaptation modules (36% in plasmids vs. 88% in chromosomes; Supplementary Figure S2), yet no significant difference in the array sizes of *cas*-associated CRISPRs. Although the reasons for the lack of adaptation modules are poorly understood, it is a characteristic feature of other MGE-encoded CRISPR-Cas systems (e.g., carried by phages and transposons) that is thought to be compensated via *in trans* use of chromosomally-encoded adaptation machinery (Peters *et al*., 2017; Faure *et al*., 2019; Pinilla-Redondo *et al*., 2019; Al-Shayeb *et al*., 2020). Finally, we observed that host chromosomes tend to carry more CRISPR arrays than plasmids; 68% of chromosomes encoding CRISPR have more than 1 array, in contrast to 36% of plasmids (Supplementary Figure S1B). Together, our results underscore a pervasive acquisition of CRISPR-Cas components by plasmids and considerable differences in the composition of plasmid- and chromosome-encoded systems.

### 2. Plasmid CRISPR-Cas subtype diversity is rich and distinct from chromosomes

We then sought to investigate the diversity of CRISPR-Cas systems across plasmid genomes. Our analysis revealed a broad range of plasmid-encoded subtypes and marked differences in their abundances (Figure 1B). Except for type VI, representatives of all CRISPR-Cas types were identified in plasmids. Overall, Class 1 systems dominate the plasmid landscape (e.g. subtypes I-E, I-B, III-B, and IV-A3), whereas Class 2 systems are poorly represented, with the notable exception of subtype V-F.

Next, we explored whether the subtype distributions on plasmids differed from those found across plasmid-associated host chromosomes. Inspection of the distribution and prevalence of CRISPR-Cas subtypes on chromosomes revealed notable differences (Supplementary Figure S3 and S4). An indicator analysis (see Materials and Methods for details) showed that IV-A3, V-F, IV-B, III-B and IV-A1 are significantly enriched subtypes for plasmid genomes when comparing all plasmids and their associated host chromosomes. A direct comparison, including only plasmid-chromosome pairs where both have CRISPR-Cas components, showed that IV-A3 is enriched on plasmids and I-D, V-J, and I-F are relatively more prevalent on chromosomes (Supplementary Figure S4). Furthermore, our analyses revealed that the higher abundance of orphan *cas* loci on plasmids (Figure 1A) is largely driven by the type IV-B systems which, consistent with previous reports (Faure *et al*., 2019; Pinilla-Redondo *et al*., 2019), are primarily encoded on plasmids and invariably lack CRISPR arrays (Figure 1B and Supplementary Figure S4). Although relatively infrequent, we found that some individual plasmids carry multiple CRISPR-Cas systems (44 out of 385 *cas*-containing loci) (Supplementary Figure S5). Among these, combinations involving type I were most common, primarily paired with type III, IV, and V, which may reflect functional compatibility between systems and, possibly, synergistic effects (Silas *et al*., 2017; Hoikkala *et al*., 2021).

We next examined the diversity of CRISPR-Cas systems on plasmids across taxa to determine the possible influence of host phylogeny on their prevalence and subtype distributions. In agreement with previous surveys across prokaryotic genomes (Makarova *et al*., 2015, 2020), our analysis revealed that the abundance of CRISPR-Cas is highly variable across host taxonomy (Figure 1C and Supplementary Figure S6). For instance, while their incidence on plasmids from *Rhodothermia*, *Deinococci* and *Clostridia* lies between 19 and 27%, in other taxa their incidence is very low or even zero. Strikingly, the prevalence and diversity of CRISPR-Cas subtypes on plasmids correlates poorly with their abundance across the chromosomes of plasmid-host taxa (Figure 1C), even when directly comparing the pool of plasmid-host chromosome pairs where both the plasmid and associated host chromosome carry CRISPR-Cas (Supplementary Figure S6). These results show distinct CRISPR-Cas compositions for plasmids and their associated host chromosomes, a pattern that likely results from the genetic autonomy of plasmids.

It is noteworthy that most available sequenced plasmids are harbored by members of Gammaproteobacteria, Bacilli, and Alphaproteobacteria (Figure 1C), which together represents 84% of all plasmids with a known host. It is therefore important to consider our results in light of this strong inherent database bias, which results from traditionally higher sampling and sequencing rates of cultivable and clinically relevant microbes (Smillie *et al*., 2010; Shintani, Sanchez and Kimbara, 2015). Consequently, given the comparatively rare occurrence of plasmid-encoded CRISPR-Cas in these dominant taxa (Figure 1C), the calculated averaged prevalence for all plasmid-encoded CRISPR-Cas systems (~3%) is predicted to be an underestimate of their true representation across environments. Taken together, our results indicate that plasmid-encoded CRISPR-Cas loci are frequent in nature and do not simply mirror those found in their host chromosomes, thereby highlighting the influence of distinct selective pressures that promote the recruitment and retention of specific subtypes on plasmids versus chromosomes.

### 3. Plasmids contribute to the horizontal dissemination of CRISPR-Cas

The recently proposed bacterial pan-immune model is based on the idea that defense systems are frequently lost and acquired by community members through HGT (Bernheim and Sorek, 2020). Therefore, we investigated whether there is a link between plasmid conjugative transmissibility and CRISPR-Cas presence. We specifically focused on proteobacterial plasmids, since high confidence predictions for conjugative transmissibility are limited to this phylum (Smillie *et al*., 2010; Shintani, Sanchez and Kimbara, 2015) and because proteobacterial plasmids dominate the dataset (62% of all non-redundant plasmid genomes).

We detected an enrichment of conjugative transfer functions within plasmids carrying CRISPR-Cas components (over 47%: average of complete systems and orphan loci; Figure 2A), a higher proportion than for plasmids not encoding CRISPR or *cas* (~36%; Fisher’s exact test: p-value = 5.9e-05; odds-ratio = 2.23). These results support the notion that conjugative plasmids facilitate HGT of CRISPR-Cas systems in the environment and, given the remarkably broad transfer ranges of some proteobacterial plasmids (e.g. IncQ, IncP, IncH and IncN) (Suzuki *et al*., 2010; Jain and Srivastava, 2013; Klümper *et al*., 2015; R. Pinilla-Redondo *et al*., 2020), possibly also across distantly related taxa. Less is known about plasmid-transfer modes outside Proteobacteria and their impact on gene exchange networks (Smillie *et al*., 2010; Shintani, Sanchez and Kimbara, 2015). For instance, many plasmids in Gram-positive bacteria transfer via conjugation but their transfer machinery is poorly characterized, thus rendering mobility predictions based on sequence data unreliable (Smillie *et al*., 2010; Garcillán-Barcia, Alvarado and de la Cruz, 2011) and highlighting that conjugative plasmids are likely underestimated in our database. Moreover, it is expected that many non-conjugative plasmids transfer horizontally through alternative mechanisms, e.g., via transformation (Lorenz and Wackernagel, 1994), mobilization (Ramsay and Firth, 2017), transduction (Ammann *et al*., 2008; Watson, Staals and Fineran, 2018), and outer membrane vesicles (Bitto *et al*., 2017). Therefore, our results underpin the idea that plasmids are major contributors to the active dissemination of CRISPR-Cas systems across microbiomes.

**Figure 2.**
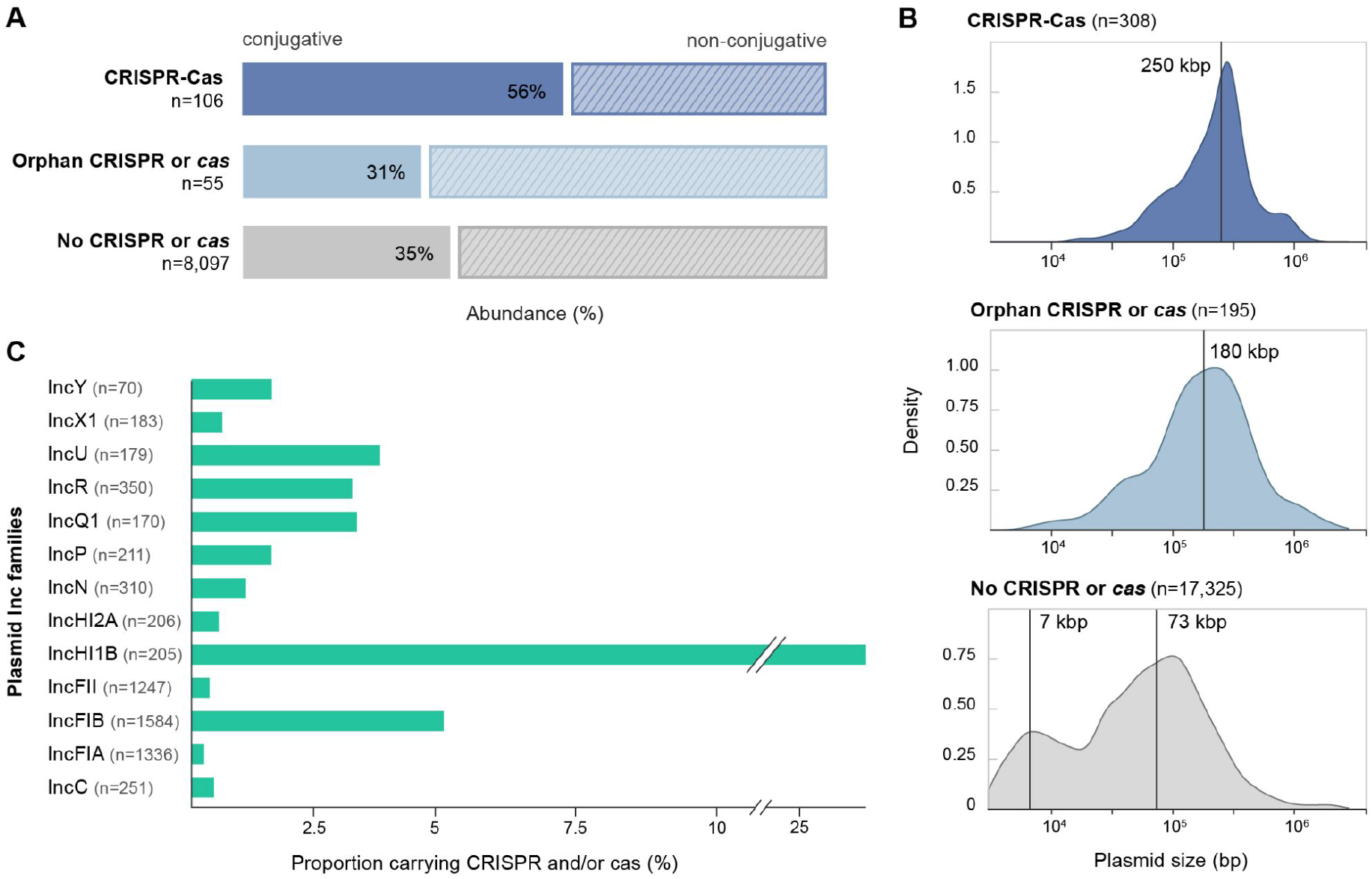
Features of plasmids encoding CRISPR-Cas components. **(A)** Mobility predictions for the collection of non-redundant proteobacterial plasmids analysed in this study, presented according to their CRISPR-Cas contents: complete CRISPR-Cas loci, orphan CRISPRs or *cas*, and no CRISPR or *cas*. **(B)** Size distributions for the collection of plasmid genomes carrying complete CRISPR-Cas loci, orphan arrays, solo *cas* operons and no CRISPR or *cas* genes. Vertical lines indicate median plasmid size for the unimodal distributions and estimated means from a 2-component gaussian mixture model for the bimodal distribution. Densities are computed with default parameters in base R. **(C)** Distribution of plasmid incompatibility groups within the Inc-typeable fraction of the complete plasmid dataset and relative abundance of the subset encoding CRISPR-Cas loci. Single plasmids can belong to more than one Inc group. Only Inc groups containing more than 10 plasmids are shown.

### 4. CRISPR-Cas systems are enriched on plasmids of larger sizes

We then sought to examine other biological characteristics of the plasmids and searched especially for common or distinctive patterns shared by CRISPR-Cas-encoding plasmids. We focused on exploring the link between plasmid genome size and the presence of CRISPR-Cas modules. In contrast to the collection of non-CRISPR-Cas-encoding plasmids—which displayed the previously reported bimodal size distribution (Smillie *et al*., 2010; Shintani, Sanchez and Kimbara, 2015)—plasmids carrying CRISPR-Cas components exhibited unimodal distributions, with the peak shifted towards larger genome sizes (180-250 Kb on average) (Figure 2B).

Given the relatively large sizes of CRISPR-Cas systems, a bias towards larger genomes is unsurprising and possibly stems from size-related constraints associated with certain plasmid life history strategies. Larger plasmids allocate considerable portions of their genomes to transfer, stabilization and accessory modules that enhance their persistence (Norman, Hansen and Sørensen, 2009). This is congruent with the observed enrichment of CRISPR-Cas systems on conjugative plasmids (Figure 2A), which are known to be relatively large and show a unimodal size distribution centered around 250 Kb (Smillie *et al*., 2010). Similar genomic streamlining dynamics appear to extend to other MGEs, including phages, where complete CRISPR-Cas systems have been reported in huge phages (>500 kb) (Al-Shayeb *et al*., 2020) but rarely occur in the more common, smaller-sized (pro)viral genomes (Faure *et al*., 2019; Bernheim *et al*., 2020). In conclusion, our data show that CRISPR-Cas systems are important components of many plasmid accessory repertoires, and are more frequently associated with plasmids of larger sizes.

### 5. Highly uneven distribution of CRISPR-Cas across plasmid Incompatibility groups

Next, we examined whether CRISPR-Cas systems in plasmids have short-lived associations or whether we could identify signs of retention by specific plasmid lineages. To this end, a common plasmid classification scheme types plasmids into incompatibility (Inc) groups and is deeply rooted in plasmid eco-evolutionary dynamics, i.e. based on the observation that plasmids sharing replication or partitioning components cannot stably propagate within a given cell host lineage (Novick, 1987). We therefore investigated the distribution and prevalence of CRISPR-Cas-containing plasmids across the Inc-typeable fraction of non-redundant plasmids, which corresponds to 29% of all plasmids (98% of which have a host belonging to Proteobacteria) (Figure 2C).

Overall, we found that only a reduced number of Inc types (15/50) include plasmids carrying CRISPR-Cas (Figure 2C and Supplementary Figure S7). Most CRISPR-Cas-encoding plasmids are distinctively concentrated within specific Inc families (e.g. IncH), underscoring the patchy distribution of CRISPR-Cas components across plasmids. Importantly, Inc families are used to infer a degree of genetic relatedness (phylogeny) and ecological cohesiveness, thus typically grouping plasmids that exhibit comparable backbone architecture, host range breadth, propagation mechanism, etc (Smillie *et al*., 2010; Redondo-Salvo *et al*., 2020). Therefore, our results indicate that some CRISPR-Cas systems are acquired by specific plasmid lineages (i.e., groups of plasmids sharing similar ecological strategies, niches and a related evolutionary trajectory) and are thus maintained stably through evolutionary timescales, presumably due to their adaptive benefits.

### 6. Plasmid spacer contents reveal a robust plasmid-targeting bias

We then focused on understanding the possible function(s) of plasmid-encoded CRISPR-Cas systems. CRISPR arrays are uniquely suited to provide ecological and biological insights; the origins of many spacer sequences can be backtracked, providing valuable clues about the functions of CRISPR-Cas and their selective benefits (Shah, Hansen and Garrett, 2009; Paez-Espino *et al*., 2013; Shmakov *et al*., 2017; Nicholson *et al*., 2019). It has been considered that the primary role of chromosome-encoded CRISPR-Cas systems is to protect cells against viruses (Makarova *et al*., 2015; Shmakov *et al*., 2017; Soto-Perez *et al*., 2019). This raised the question as to whether plasmid-encoded CRISPR-Cas components reinforce this function, especially given that many plasmids encode genes that enhance the fitness of their hosts against diverse environmental threats (e.g. antimicrobial resistance) (Norman, Hansen and Sørensen, 2009; Rankin, Rocha and Brown, 2011).

All spacer sequences (n=11,080) were extracted from the bacterial and archaeal plasmid-encoded CRISPR arrays and searched against comprehensive virus and plasmid sequence datasets (Material and Methods). For comparison, analogous searches were performed with the collection of spacers originating from: 1) the host chromosomes associated with the plasmids in this study (a total of 96,870 spacers), and 2) plasmid-host chromosome pairs where both the plasmid and associated host chromosome carry at least one CRISPR array (4,816 plasmid spacers and 10,315 chromosomal spacers). Only a limited fraction of spacers yielded significant matches to protospacer sequences (plasmids: 11.1%; hosts: 12.9%), consistent with previous studies (Mojica *et al*., 2009; Shah, Hansen and Garrett, 2009; Stern *et al*., 2010; Touchon and Rocha, 2010; Shmakov *et al*., 2017, 2020; Pinilla-Redondo *et al*., 2019). This is ascribed to a combination of factors, including the paucity of mobilome sequences across public databases and the high mutation rates of MGE protospacers, presumably to escape CRISPR-Cas targeting (Andersson and Banfield, 2008; Touchon and Rocha, 2010; Shmakov *et al*., 2017).

Subsequently, we examined the origins of these protospacer targets. Strikingly, a larger fraction of plasmid spacers matched sequences from other plasmids (66%), while a substantially smaller fraction matched viruses (27%) (Figure 3A). In contrast, the spacer contents originating from plasmid-host chromosomes revealed the opposite trend: a larger proportion of spacers matched viral sequences compared to plasmids (62% and 24%, respectively) (Figure 3B; Supplementary Figure S8)– consistent with a primary antiviral role of chromosomal CRISPR-Cas systems (Paez-Espino *et al*., 2013, 2015; Makarova *et al*., 2015; Shmakov *et al*., 2017; Nasko *et al*., 2019; Soto-Perez *et al*., 2019). Importantly, a more direct examination of plasmid-host chromosome pairs (limited to comparisons where both parties carry at least one CRISPR-Cas system) revealed an analogous targeting trend (Supplementary Figure S9A-B).

**Figure 3.**
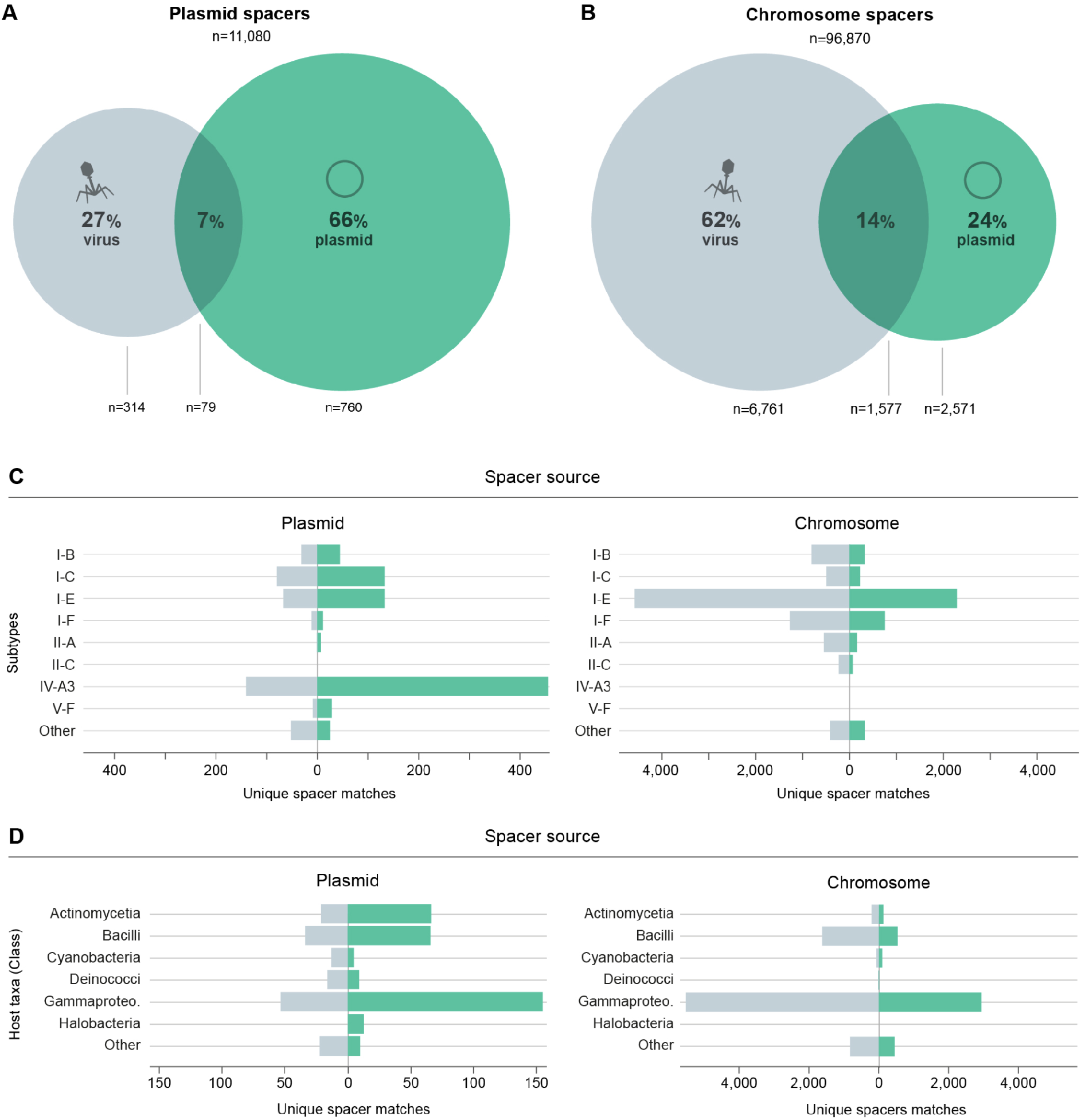
Analysis of plasmid CRISPR spacer contents reveals a plasmid targeting bias. The proportion of plasmid (**A**) and plasmid-host (**B**) chromosomal spacers matching plasmids (green) and viruses (grey). **(C)** Distribution of spacer-protospacer matches derived from plasmid (left) and plasmid-host chromosome (right) spacer contents, presented according to CRISPR-Cas subtype/variant and predicted spacer target: plasmids (green) and viruses (grey). Only the top 8 subtypes for both plasmids and chromosomes are represented, with the remaining grouped in “Other”. **(D)** Distribution of spacer-protospacer matches derived from plasmid (left) and host chromosome (right) spacer contents, broken down by host taxa and predicted protospacer origin: plasmids (green) and viruses (grey). Only the top 6 classes for both plasmids and chromosomes are represented, with the remaining grouped in “Other”.

The abundance of plasmid spacers targeting other plasmids raised the question of whether the reported plasmid-targeting preference of type IV CRISPR-Cas systems (Pinilla-Redondo *et al*., 2019) could be driving this trend, especially given the abundance of type IV spacers within our dataset (12% of plasmid spacers, yielding 48% of the spacers with any match) (Supplementary Figure S10A). However, we found that the plasmid-targeting bias also held true for the majority of other plasmid-encoded CRISPR-Cas subtypes/variants (Figure 3C and Supplementary Figure S9C). In contrast, chromosomal spacers maintained a virus-targeting preference, regardless of CRISPR-Cas subtype (Figure 3C and Supplementary Figure 9C). Furthermore, we found that the plasmid-to-plasmid vs. chromosome-to-virus targeting patterns are maintained across the different taxa, implying the existence of a robust biological underpinning of this phenomenon (Figure 3D and Supplementary Figure S9D). Nevertheless, the plasmid-encoded CRISPRs from certain underrepresented taxa (Figure 3D and Supplementary Figure S9D) appear to be enriched with virus-targeting spacers (e.g. Rhodothernia and Cyanobacteria), suggesting that enhancement of antiviral host immunity could still be an important evolved strategy for some groups of plasmids.

### 7. A reticulated web of CRISPR-based plasmid-plasmid targeting

The identification of extensive plasmid-plasmid targeting provides a practical framework for investigating plasmid eco-evolutionary dynamics and offers a unique opportunity to gain insights into HGT routes. This prompted us to build a global network of plasmid-plasmid interactions based on the linkage information provided by the CRISPR-targeting data (Figure 4A). The corresponding directed graph consists of de-replicated plasmid genomes (nodes), connected by the predicted spacer-protospacer matches (edges). Overall, our analyses revealed a network with a pronounced modular structure, where a reduced number of densely connected clusters accrue the majority of plasmids, and links between clusters are very sparse. A highly visible trend across the targeting network is the clustering of plasmids according to host taxonomy, with the two largest clusters consisting of plasmids from either Enterobacteriales or Bacillales. However, generalisations based on such a trend should be made with caution and viewed in the context of the historical sequencing bias towards plasmids from cultivable and/or clinical strains. For example, inferring that plasmid targeting is a distinctive phenomenon among Enterobacteriales or Bacillales plasmids could be an inaccurate assumption, since plasmids carrying CRISPR-Cas are relatively rare in these taxa (Figure 1C), despite their sequences comprising the overwhelming majority of sequenced plasmids (Figure 1C). As more accurate sampling and sequence representation of plasmid diversity becomes available, a more clear understanding of plasmid-plasmid targeting will emerge.

**Figure 4.**
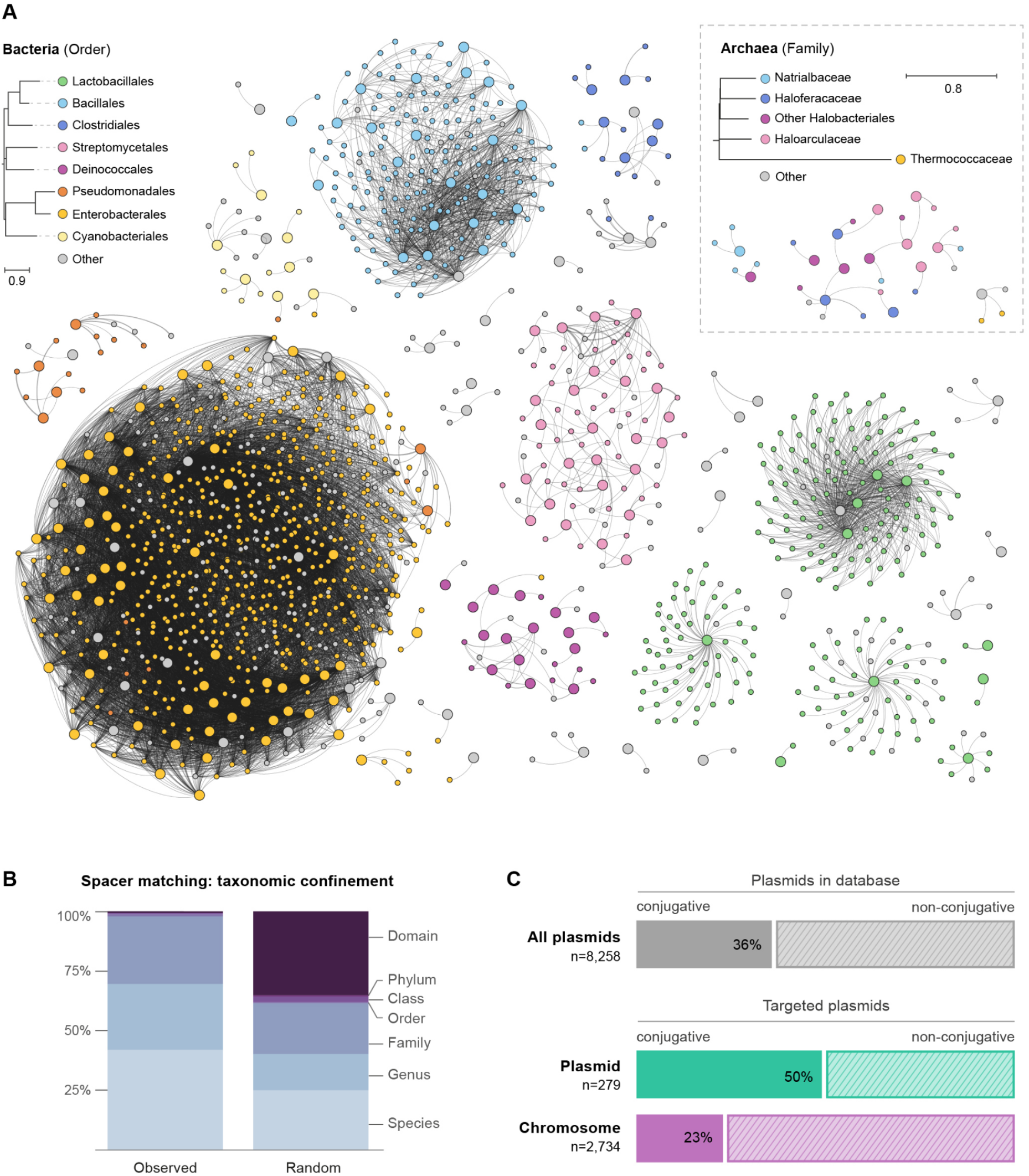
Plasmid-plasmid CRISPR-Cas targeting network reveals a clustering organisation showing high concordance with host taxonomy. **(A)** Global representation of plasmid-plasmid targeting network colored at the host order level. Nodes correspond to individual plasmids and edges represent CRISPR-Cas targeting, based on predicted spacer-protospacer matches. Large and small nodes indicate the presence or absence of CRISPR-Cas in the plasmid, respectively. Edge thickness is proportional to the number of spacer-protospacer matches between plasmid pairs. The phylogeny in the legend is based on the median cophenetic distance from the GTDB whole-genome phylogenies, with the tree inferred by neighbor-joining. “Other” indicates plasmids without a known host, host with a different taxonomy than those displayed, or with a host with unspecific taxonomy. An expanded view of plasmid-plasmid targeting within the class Gammaproteobacteria can be found in Supplementary Figure S11. **(B)** Taxonomic confinement of plasmid-plasmid CRISPR-Cas targeting predictions. The percentage distribution of spacer-protospacer matches circumscribed within the different plasmid-host taxonomic levels (observed) is compared to their distribution when the taxonomic labels are randomly permuted among the set of targeting plasmid-plasmid pairs (random). **(C)** Proportion of conjugative plasmids targeted by plasmid and plasmid-host chromosomal spacers. The incidence of conjugative plasmids across PLSDB is shown below. This assessment is restricted to Proteobacterial plasmids.

Notably, the clustering analysis demonstrates a pronounced inverse relationship between the number of plasmid connections and the phylogenetic hierarchy of the cognate bacterial hosts. Whereas targeting between plasmids within a single species, genus and family account for the bulk of all predictions (~42%, 28% and 28%, respectively), matches confined to higher taxa comprise less than 2% (Figure 4B). Indeed, a closer examination of the plasmid-plasmid targeting network in Gammaproteobacteria revealed abundant links between plasmids from different genera (Supplementary Figure S11). These results underscore that taxonomic boundaries represent a major hurdle for plasmid dissemination. Indeed, although some plasmids are able to transfer between distantly-related taxa, their long-term evolutionary host range is primarily constrained to a narrower group of phylogenetically related hosts (Acman *et al*., 2020; Redondo-Salvo *et al*., 2020). Furthermore, acquisition of spacers from plasmids sharing a similar host range is expected to be more frequent due to the conceivably higher rates of encounters within cells. From a CRISPR-targeting standpoint, spacer retention is also likely influenced by the selective advantage they can provide in plasmid-plasmid competition dynamics.

Given the self-transmissible properties of conjugative plasmids, we wondered whether their effective spread through bacterial populations could render them common targets for CRISPR-Cas compared to non-conjugative plasmids. In support of this, we observed an over-representation of plasmid spacers predicted to target conjugative plasmids (Figure 4C). This may indicate that conjugative invasion is detrimental to plasmids already established in a cell. This is consistent with previous reports of plasmid-encoded mechanisms specifically directed towards preventing the entry of conjugative plasmids (e.g., fertility inhibition strategies and entry exclusion systems) (Getino and de la Cruz, 2018). Interestingly, we found that chromosomal spacers showed a relative underrepresentation of spacers targeting conjugative plasmids, suggesting that this type of plasmids may be less detrimental to bacteria, possibly owing to the fitness benefits associated with the adaptive gene cargos that they frequently carry (Norman, Hansen and Sørensen, 2009; Smillie *et al*., 2010).

## DISCUSSION

The study of CRISPR-Cas biology has primarily focused on chromosomally-encoded systems and their adaptive antiviral functions in bacteria and archaea. While recent work has started to uncover the common association of CRISPR-Cas systems with diverse MGEs and the importance of this phenomenon for CRISPR-Cas ecology and evolution (Faure *et al*., 2019), their recruitment by plasmids has remained largely unexplored. Here, we present the first comprehensive analysis of CRISPR-Cas systems across the largest curated dataset of wildtype bacterial and archeal plasmids. We show that CRISPR-Cas components are pervasive accessory components of many plasmids and span a broad diversity of systems, including subtype representatives covering five out of the six known types. Interestingly, we found that certain plasmids carry multiple CRISPR-Cas systems (Figure 5A). The incidence of plasmid-encoded systems is highly uneven across taxa—ranging from 0 to 30%, but averaging at ~3%—and the subtype diversity does not simply reflect the CRISPR-Cas contents found in the chromosomes of their host. Our results thus underscore the genetic independence of plasmids and the influence of distinct evolutionary pressures in the acquisition and retention of CRISPR-Cas on plasmids versus their associated host chromosomes.

**Figure 5.**
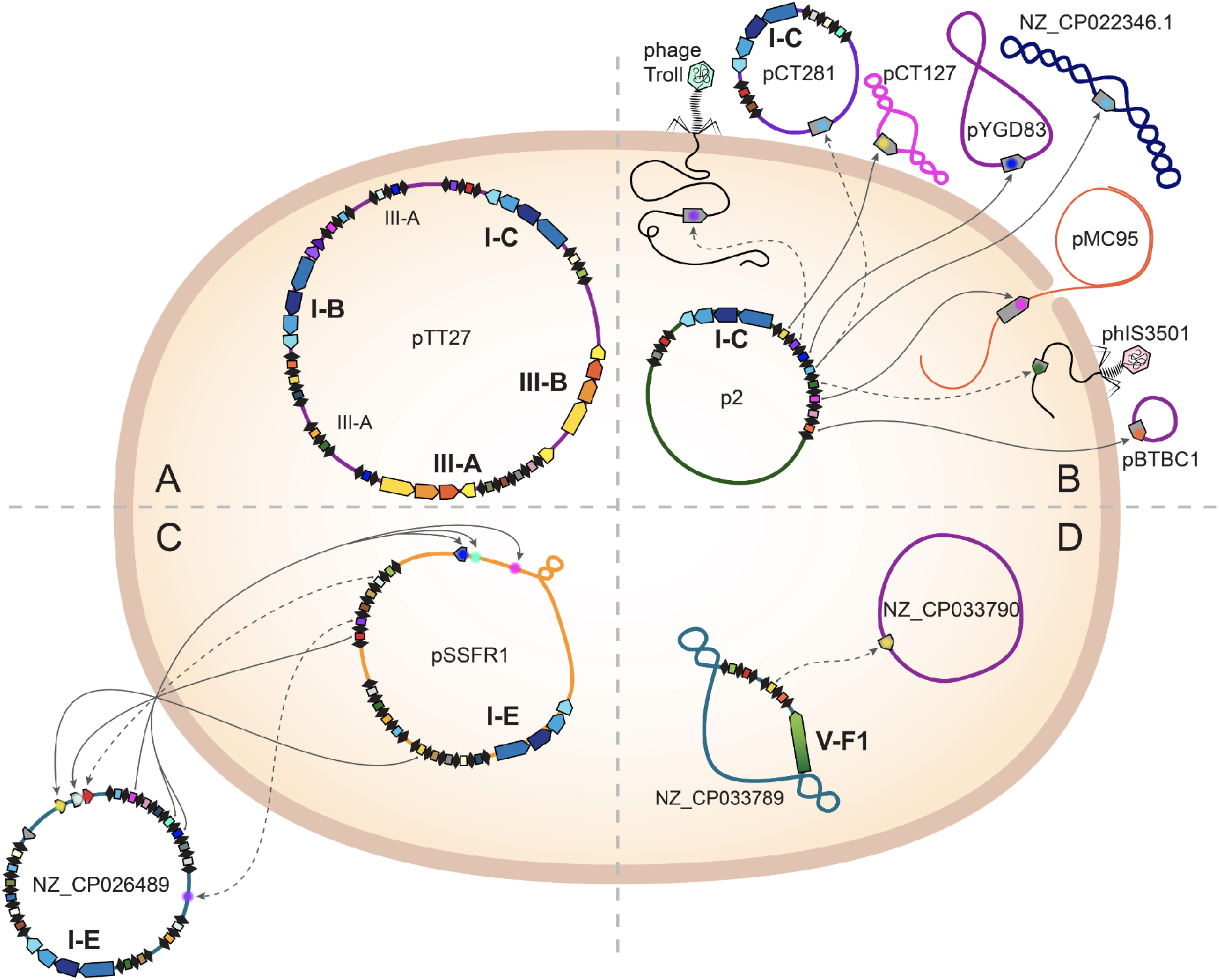
Plasmid-encoded CRISPR-Cas systems mediate complex plasmid-MGE interactions. Diagram of representative plasmid-encoded CRISPR-Cas systems and analysis of the predicted targeting dynamics. *Cas* operons are color-coded according to their classification (type level): type I in blue, type III in orange, and type V in green; CRISPR arrays are depicted as serial black diamonds (repeats) interspaced by colored rectangles (spacers). Arrows indicate identified spacer-protospacer matches, where each protospacer is represented in a different color. Solid lines denote spacer targets with no or one mismatch and dashed lines denote two to four mismatches. The name or accession number of plasmids and phages are indicated. **(A)** Plasmid carrying multiple different CRISPR-Cas systems. **(B)** Plasmid predicted to target diverse MGEs (7 plasmids and 2 phages). **(C)** Plasmids predicted to target each other (cross-targeting). **(D)** Plasmid targeting another plasmid residing in the same cell.

Intriguingly, putatively incomplete loci were more abundant on plasmids than chromosomes, although less abundant than previously reported (Bernheim *et al*., 2020). It has been suggested that orphan CRISPR arrays and *cas* loci may be remnants of decaying CRISPR-Cas systems (Makarova *et al*., 2015). Their relatively higher occurrence on plasmids could indicate that CRISPR-Cas systems erode faster on plasmids, or that orphan components are recruited and/or selectively maintained to perform important, but as yet unknown, biological functions. Orphan CRISPR arrays could, for instance, employ host Cas machinery *in trans* (Almendros *et al*., 2016; Deecker and Ensminger, 2020) or facilitate plasmid chromosome integration via recombination between plasmid and host-encoded CRISPRs (Varble *et al*., 2019, 2020). On the other hand, the higher proportion of orphan components may be an artefact of CRISPR-Cas prediction tools unable to detect a conceivably greater diversity of uncharted (sub)types across plasmids. Indeed, novel subtypes have recently been identified on diverse MGEs (Peters *et al*., 2017; Faure *et al*., 2019; Pinilla-Redondo *et al*., 2019; Al-Shayeb *et al*., 2020).

The observed enrichment of conjugative functions across CRISPR-Cas-encoding plasmids, together with the expected underestimation of transmissible plasmids in our database (i.e., due to unreliable bioinformatic prediction methods and unknown plasmid mobility mechanisms), suggest an active contribution of plasmids to the conspicuous dissemination of CRISPR-Cas systems across microbiomes. These results are in agreement with the proposed bacterial pan-immune concept, where defense systems are continually lost and (re)gained by bacteria through HGT mechanisms (Bernheim and Sorek, 2020), and further consistent with the common observation of restriction-modification and toxin-antitoxin systems on plasmids (Van Melderen, 2010; Rego, Bestor and Rosa, 2011).

Notably, we found that plasmid-encoded CRISPR arrays tend to carry a larger fraction of spacers predicted to target other plasmids, while plasmid-host chromosome-encoded systems show the commonly observed targeting bias towards viruses. This contrasting targeting preference was consistently observed across taxa and the different CRISPR-Cas subtypes, indicating that plasmids may primarily exploit CRISPR-Cas systems to target other plasmids, and thus likely play a less dominant role in host protection against viral predators (Figure 5B). These observations extend the hypothesis that the main function of plasmid-encoded type IV CRISPR-Cas systems is to eliminate plasmid competitors (Crowley *et al*., 2019; Pinilla-Redondo *et al*., 2019). Interestingly, we found a number of cases of plasmid cross-targeting pairs (26 in total, 4 de-replicated), where CRISPR-Cas-encoding plasmids are predicted to target each other upon crossing paths in a host cell (Figure 5C). We also found 29 examples of plasmids predicted to target other plasmids within the same cell (Figure 5D), which could indicate the presence of counter-defense strategies to avoid targeting, such as plasmid-encoded anti-CRISPRs (Acrs) (Mahendra *et al*., 2020; Rafael Pinilla-Redondo *et al*., 2020). Although we failed to identify known Acrs across the co-residing targeted plasmids, recent work describes an analogous co-evolutionary arms race between a conjugative island-encoded I-C CRISPR-Cas system and diverse MGE-encoded Acrs in *Pseudomonas aeruginosa* (Leon *et al*., 2021).

Together, our results are consistent with previous reports of the co-option of CRISPR-Cas systems, or components thereof, by different MGEs for waging inter-MGE conflicts. For example, the ICP1 *Vibrio cholerae* phage encodes a I-F CRISPR-Cas system to restrict the phage satellite PLE, a MGE that parasitizes ICP1 (Seed *et al*., 2013; McKitterick *et al*., 2019). Additionally, some giant phages and other viruses carry either complete CRISPR-Cas systems or “mini-arrays’’ that might contribute to inter-viral conflicts (Minot *et al*., 2011; Faure *et al*., 2019; Medvedeva *et al*., 2019; Al-Shayeb *et al*., 2020). Our findings thus support the “guns for hire” concept (Koonin *et al*., 2020), whereby CRISPR-Cas systems are continually repurposed by different genetic entities. Because similar entities are expected to compete more strongly due to niche overlap (e.g. space and resources), it is not surprising to observe CRISPR-Cas driven inter-viral and inter-plasmid conflicts. Moreover, the higher proportions of virus-derived chromosomal spacers found here, and earlier, illustrate how viruses exert a stronger selection on hosts than plasmids do. Indeed, while viruses often kill their host cell, plasmids tend to only affect fitness - and can be beneficial under certain conditions. Together, these results suggest that retention of CRISPR spacer content is primarily shaped by the selective advantage single spacers confer on the genetic entities carrying them and to a lesser extent by any possible biases inherent to the spacer acquisition and targeting mechanisms.

More broadly, the implications of our findings have practical applications beyond CRISPR-Cas biology. Plasmid sequences may hide an uncharted diversity of CRISPR-Cas systems with promising biotechnological applications, e.g. in genome engineering. Furthermore, plasmid-derived CRISPRs can be exploited to determine information about a plasmid’s direct relationships with other elements across evolutionary timescales. When available, spacer-protospacer match prediction data could comprise an added layer of information during retrospective plasmid host-range inference analyses, similar to how chromosomal CRISPR contents are leveraged for bioinformatic deconvolution of virus-host associations (Anderson, Brazelton and Baross, 2011; Sanguino *et al*., 2015; Hidalgo-Cantabrana, Sanozky-Dawes and Barrangou, 2018; Shmakov *et al*., 2020; Dion *et al*., 2021). Furthermore, the distinctive spacer acquisition bias at the leader end of most CRISPR arrays (Andersson and Banfield, 2008; Jackson *et al*., 2017) suggests a promising resource for extracting chronological information about plasmid dissemination routes. Such analyses may become particularly valuable in the study of clinically relevant plasmids (e.g. those carrying antibiotic resistance or virulence determinants), for which plasmid typing and epidemiological tracking are crucial but currently difficult to infer through sequence analyses alone (Suzuki *et al*., 2010; Sen *et al*., 2013; Pinilla-Redondo *et al*., 2018; Redondo-Salvo *et al*., 2020).

Overall, CRISPR-Cas systems constitute powerful barriers against MGE-mediated HGT in microbial communities. While the investigation of CRISPR-Cas biology has focused on chromosomally-encoded systems, our work uncovers their pervasive association with plasmids across a broad phylogenetic breadth, where they appear to play a major role in mediating plasmid-plasmid conflicts. We anticipate that MGE-MGE warfare likely constitutes an important, yet largely overlooked, factor influencing the dynamics of gene flow across microbiomes.

## MATERIAL AND METHODS

### Software and code availability

Scripts for downloading data and reproducing all analyses are available at https://github.com/Russel88/CRISPRCas_on_Plasmids. Analyses were made with a combination of shell, python 3, and R 3.6.3 scripting. Plots were made with ggplot2, heatmaps with pheatmap, phylogenetic trees with iTOL (Letunic and Bork, 2021), and networks with gephi (Bastian, Heymann and Jacomy, 2009).

### Dataset construction

A total of 27,939 complete bacterial plasmid sequences were downloaded from PLSDB 2020_11_19 (https://ccb-microbe.cs.uni-saarland.de/plsdb) (Galata *et al*., 2019), together with their associated metadata (Galata *et al*., 2019). A total of 253 manually curated archaeal plasmids were downloaded from NCBI RefSeq on January 6th 2020. Plasmid-host chromosome associations were determined through the NCBI assembly information, for which only sequences annotated as “chromosome” were included as host sequences. Using this approach, we were able to assign a host for 21,974 of the plasmids. The number of archaeal plasmids selected is relatively low because few archaeal plasmids have been characterised and sequenced (Lipps, 2007). We used GTDBtk v1.4.1 (Chaumeil *et al*., 2019) to re-annotate the taxonomy of the host of each plasmid in a common phylogenomic framework. To filter out redundant plasmids, they were de-replicated using dRep version 3.1.0 (Olm *et al*., 2017) with the following parameters: 90% ANI cut-off for primary clustering, 95% ANI cut-off for secondary clustering and a total coverage of 90%, with fastANI (Marçais *et al*., 2018) as secondary clustering algorithm. Size was the only criterion used to choose the plasmid to include in each cluster, such that the largest plasmid (or random among these given ties) was picked among the clustered plasmids. Dereplication resulted in a total of 17,828 plasmids, out of which 13,265 could be associated with known prokaryotic hosts.

### Identification of CRISPR loci

Detection of CRISPR arrays was carried out by using CRISPRCasFinder 4.2.17 (Couvin *et al*., 2018), coupled to an optimized algorithm for false-positive array removal (Supplementary Figure S12) and an additional analysis for finding CRISPR loci that are commonly missed by this algorithm. Briefly, high confidence arrays predicted by CRISPRCasFinder (evidence level 4) were automatically kept. The remaining arrays were binned into a “quarantine list” if they were found to clear a series of conservative manually-curated parameter cutoffs: 1) calculated average CRISPR repeat conservation across the array > 70%, 2) spacer conservation < 50%, 3) standard error of the mean of the array’s spacer lengths < 3, and 4) array does not overlap with an open reading frame (ORF) with a prediction confidence of at least 90% (Hyatt *et al*., 2010). Putative arrays from the quarantined list were rescued for further analysis if they were found within 1 Kb to a predicted *cas* gene or matched (95% coverage and 95% identity) with any previously defined high confidence CRISPR repeat: CRISPRCasFinder evidence level 4 or archived in CRISPRCasdb (Pourcel *et al*., 2020). This upgrade reduced the rate of detection of false positive CRISPRs, most of which constitute short repetitive genomic regions that are erroneously selected by CRISPRCasFinder (Zhang and Ye, 2017), and which are more common on plasmids (e.g. iterons and tandem transposon-associated repeats) (Chattoraj, 2000; Giraldo and Fernández-Tresguerres, 2004; Oliveira *et al*., 2010). High confidence CRISPR repeats (see above) were then BLASTed (task: blastn-short, 95% coverage and 95% identity) to a database in which the CRISPR loci that were already detected were masked and any matches within 100 bp were clustered into arrays. Arrays with less than 3 repeats were excluded from all analyses.

### Identification and typing of *cas* loci

The prediction and classification (at the subtype or variant level) of *cas* operons was carried out by CRISPRCasTyper 1.2.4 (https://github.com/Russel88/CRISPRCasTyper) (Russel *et al*., 2020). CRISPR arrays closer than 10 Kbp to the nearest *cas* operon were considered to be linked; the 10 Kbp cutoff was based on an analysis of the distribution of distances of CRISPR arrays to the closest *cas* operon (Supplementary Figure S13). Furthermore, we used CRISPR-repeat similarity information to type arrays that were not found linked to *cas* operons. These distant arrays (>10 Kbp from the nearest *cas* operon) were considered associated with a *cas* operon if the direct repeat sequence was at least 85% identical to the direct repeat sequence of an array adjacent to that *cas* operon (Supplementary Figure S14). When possible, CRISPR-Cas systems annotated as “Ambiguous” were manually subtyped.

### Indicator analysis

Enrichment of certain CRISPR-Cas subtypes on either plasmids or host chromosomes was investigated with an indicator species analysis, using the indicspecies R package. For the comparison between all plasmids and chromosomes the IndVal.g statistic was used, which controls for difference in group sizes. For the direct comparison between plasmids and hosts chromosomes, where both carry CRISPR-Cas, the IndVal statistic was used. Statistical significance was determined by permutation (n=9999) and a Bonferroni adjusted p-value threshold of 0.05 was used.

### Plasmid conjugative transfer and incompatibility group prediction

The conjugative transfer functions and incompatibility (Inc) typing of all plasmids in PLSDB was predicted with MOB-suite v3.0.1 using *mob_typer* function (Robertson and Nash, 2018) using default parameters.

### Spacer-protospacer match analysis

The genomic regions where CRISPR arrays were identified on plasmids (including CRISPR arrays with 2 repeats, which were otherwise excluded from the analyses) were masked in order to avoid false positive matches to spacers in arrays. Furthermore, for matches to plasmids only matches to high confidence ORFs were included, also to rule out any matches to possibly undetected CRISPR arrays. Spacers from orphan arrays whose consensus repeat could not be typed by repeatTyper from CRISPRCasTyper (https://typer.crispr.dk, model version 2021_03 (Russel *et al*., 2020)) were excluded from the spacer analysis to avoid any bias stemming from possible false positive arrays in this group.

Viral genomes were obtained from the IMG/VR v3 (2020-10-12_5.1, (Roux *et al*., 2021)) only including those annotated as “Reference”, which includes 39,296 viral genomes. Spacer sequences from plasmids and plasmid-associated host chromosomes were aligned against the masked dereplicated plasmid database and the virus database using FASTA 36.3.8e (Pearson, 2014). Alignments were filtered using an e-value cutoff of 0.05. To reduce redundancy bias, spacers were only counted once, no matter the absolute number of matches.

Networks were visualized in gephi with layout generated by a combination of OpenOrd and Noverlap algorithms. For calculating taxonomic confinement of spacer-protospacer matches between plasmids, each pair of plasmids connected by at least one spacer-protospacer match was counted as one matching pair. Cross-targeting plasmids were included as two separate plasmid pairs. Confinement was calculated as the number of matches found exclusively within a specific taxonomic rank, such that each plasmid-plasmid pair was only counted once. For estimating confinement of random spacer-protospacer matching, the taxonomic annotations were permuted among the plasmid-plasmid pairs with observed spacer-protospacer matches. This was repeated 100 times and the median number of matches was used as an estimate of confinement for hypothetically random matches. For estimating targeting bias towards conjugative vs. non-conjugative plasmids each unique spacer was counted with a weight of 1 with the targeting bias proportional to the number of matches to conjugative and non-conjugative plasmids, respectively. For example, a spacer matching 4 conjugative plasmids and 1 non-conjugative plasmids is counted as 0.8 for conjugative matches and 0.2 for non-conjugative matches.

## ACKNOWLEDGEMENTS

R.P-R. was financed by the Independent Research Fund Denmark, InTrans Project [#8022-00322B]. JR was supported by the Novo Nordisk Foundation. DMM was supported by a University of Otago Doctoral Scholarship. PCF was supported by the Bio-Protection Research Centre (Tertiary Education Commission). We are thankful to Marina Pinilla for her contributions in the creative design of figures. We thank members of our laboratories for helpful discussions and mimosas. We are especially thankful to Sarah Camara, Mario R. Mestre, Leah M. Smith and Nils Birkholz for valuable input and helpful discussions.

## COMPETING INTERESTS

The authors declare no conflicts of interest.

## SUPPLEMENTARY MATERIAL

**Supplementary Figure S1.**
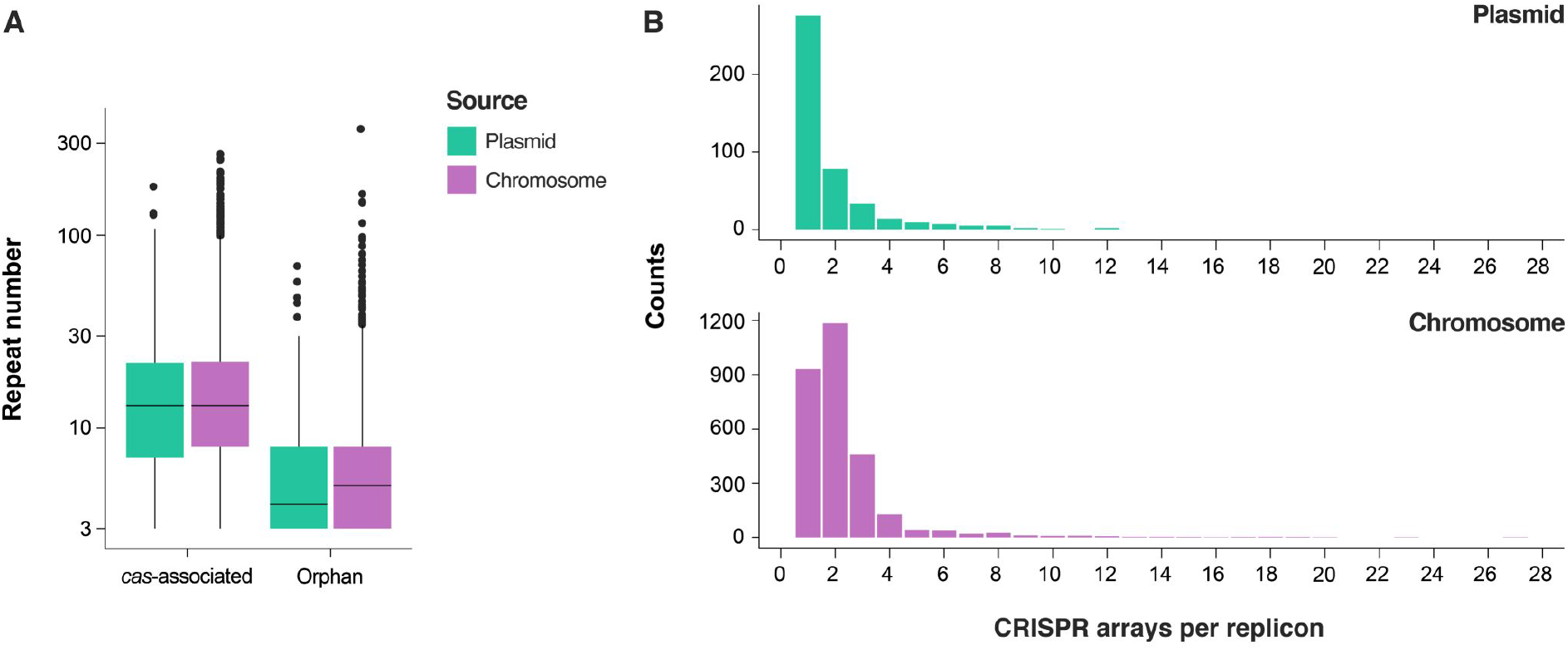
**(A)** Average array lengths of orphan and *cas*-associated CRISPRs originating from plasmids (green) and plasmid-host chromosomes (purple). Orphan CRISPRs have significantly fewer repeats (p<2e-16, negative-binomial GLM), but in each group there is no difference between chromosomal and plasmid CRISPR lengths (p=0.248), and no interaction between the two (p=0.134) **(B)** Frequency distribution of the number of CRISPR arrays per plasmid and plasmid-host chromosomes. Only replicons with at least 1 array are displayed.

**Supplementary Figure S2.**
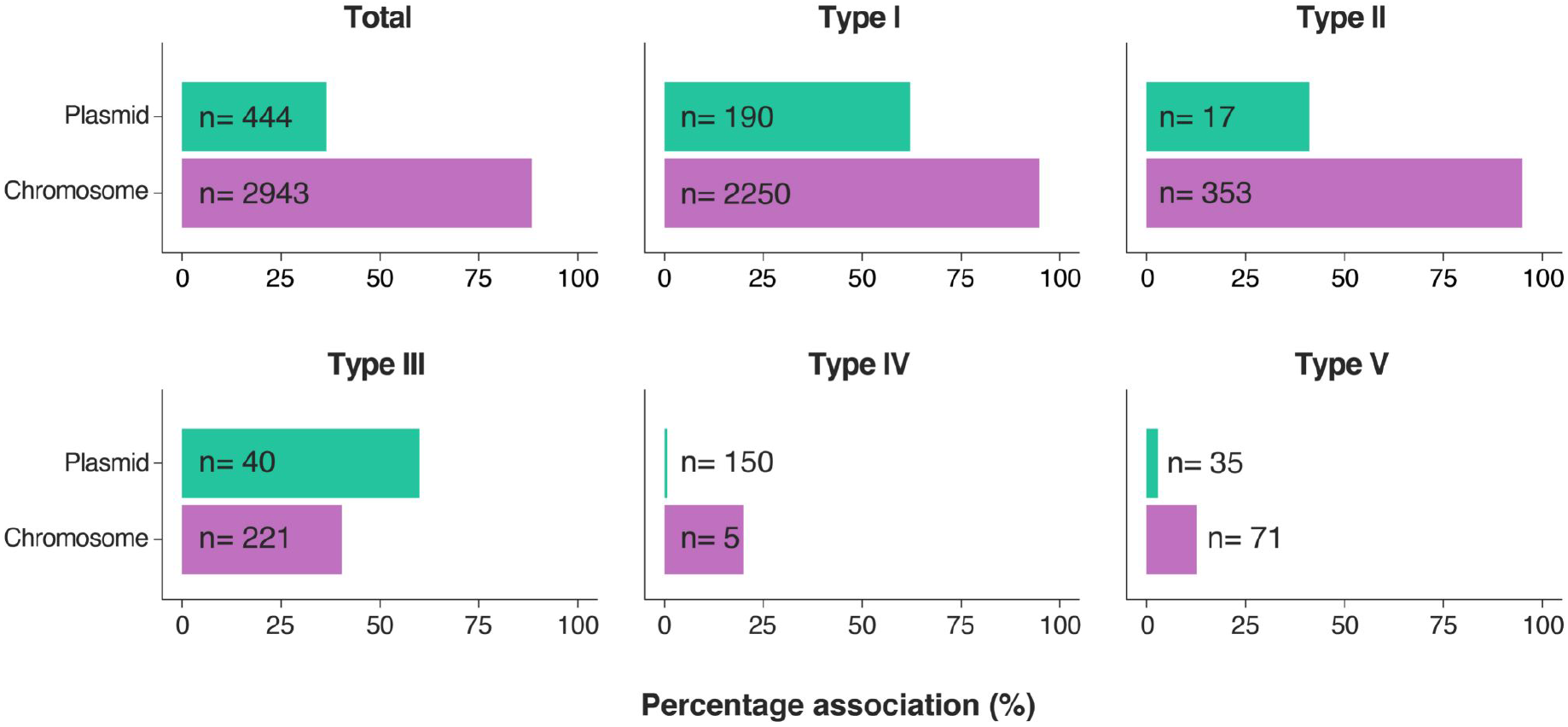
Association of adaptation modules with CRISPR-Cas systems encoded by plasmids and plasmid-host chromosomes. Percentage of CRISPR-Cas systems, separated at the type level, predicted to encode adaptation components (Cas1 and/or Cas2). The aggregate value (averaging data from all CRISPR-Cas types) is shown under “Total”.

**Supplementary Figure S3.**
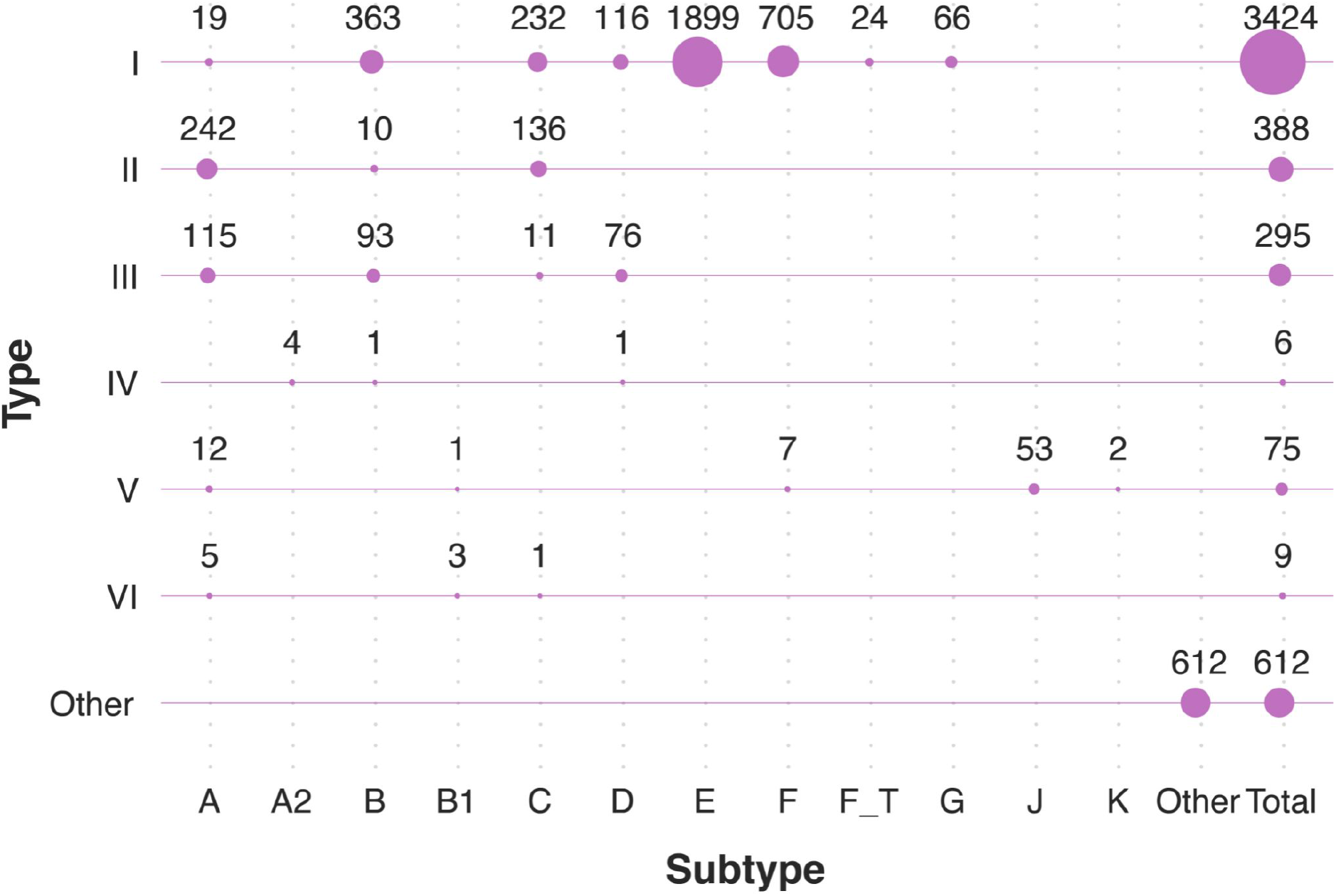
Distribution and prevalence of CRISPR-Cas subtypes across plasmid-associated host chromosomes. “Other” represents systems that could not be unambiguously assigned (e.g. multiple equally scoring subtypes, co-localised/hybrid systems, and orphan-untyped components). “I-F_T” refers to the transposon-associated subtype I-F variant and subtype IV-A is subdivided into its known variants (IV-A1 to A3). Total counts per CRISPR-Cas type are summarised on the right.

**Supplementary Figure S4.**
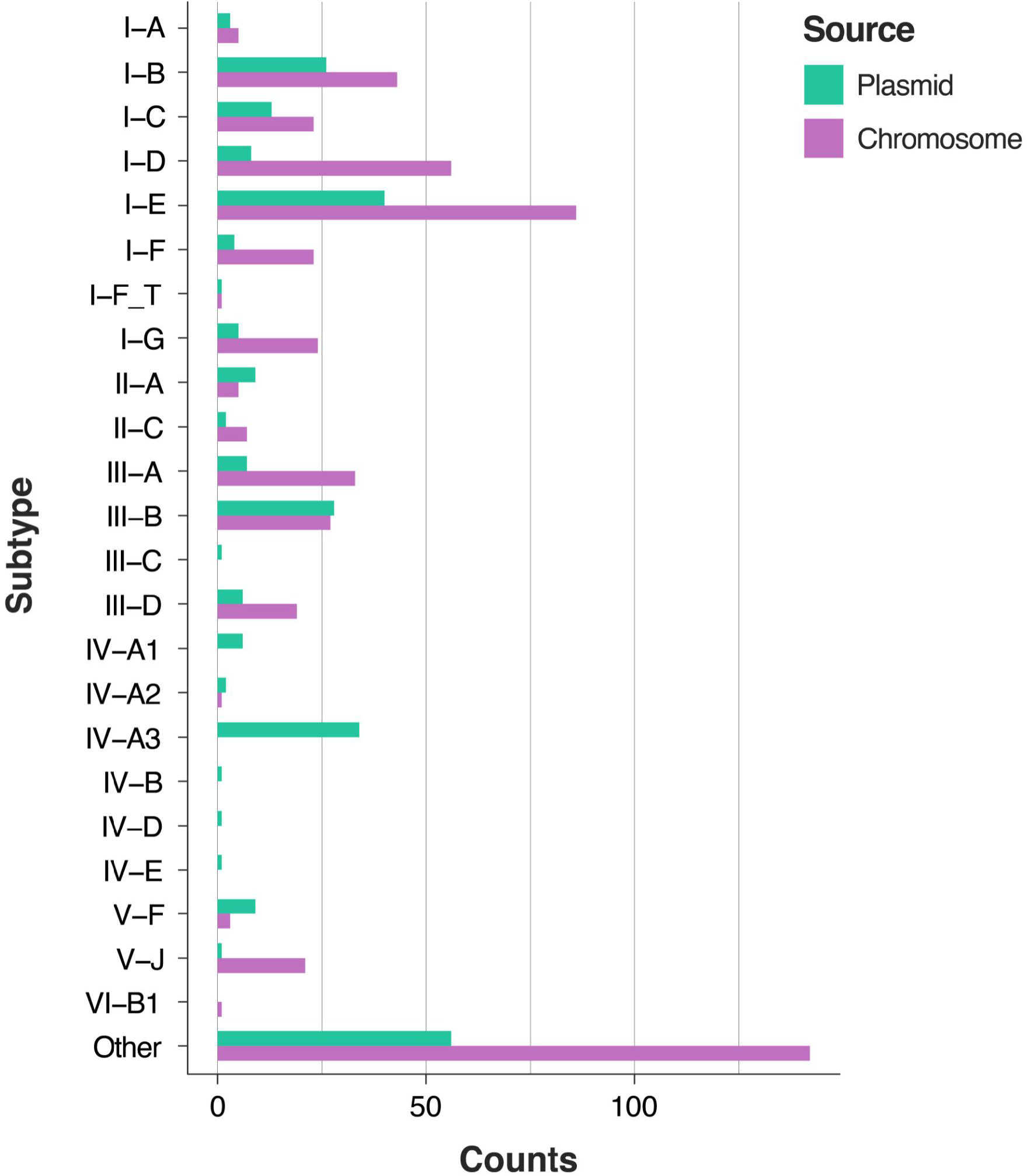
Direct comparison of the distribution and prevalence of CRISPR-Cas subtypes across plasmids and their plasmids-associated chromosomes, where the plasmid and chromosome each carry at least one CRISPR-Cas. Number of plasmid- and chromosomal systems (green and purple, respectively) are shown, broken down at the subtype level (or variant level, when possible).

**Supplementary Figure S5:**
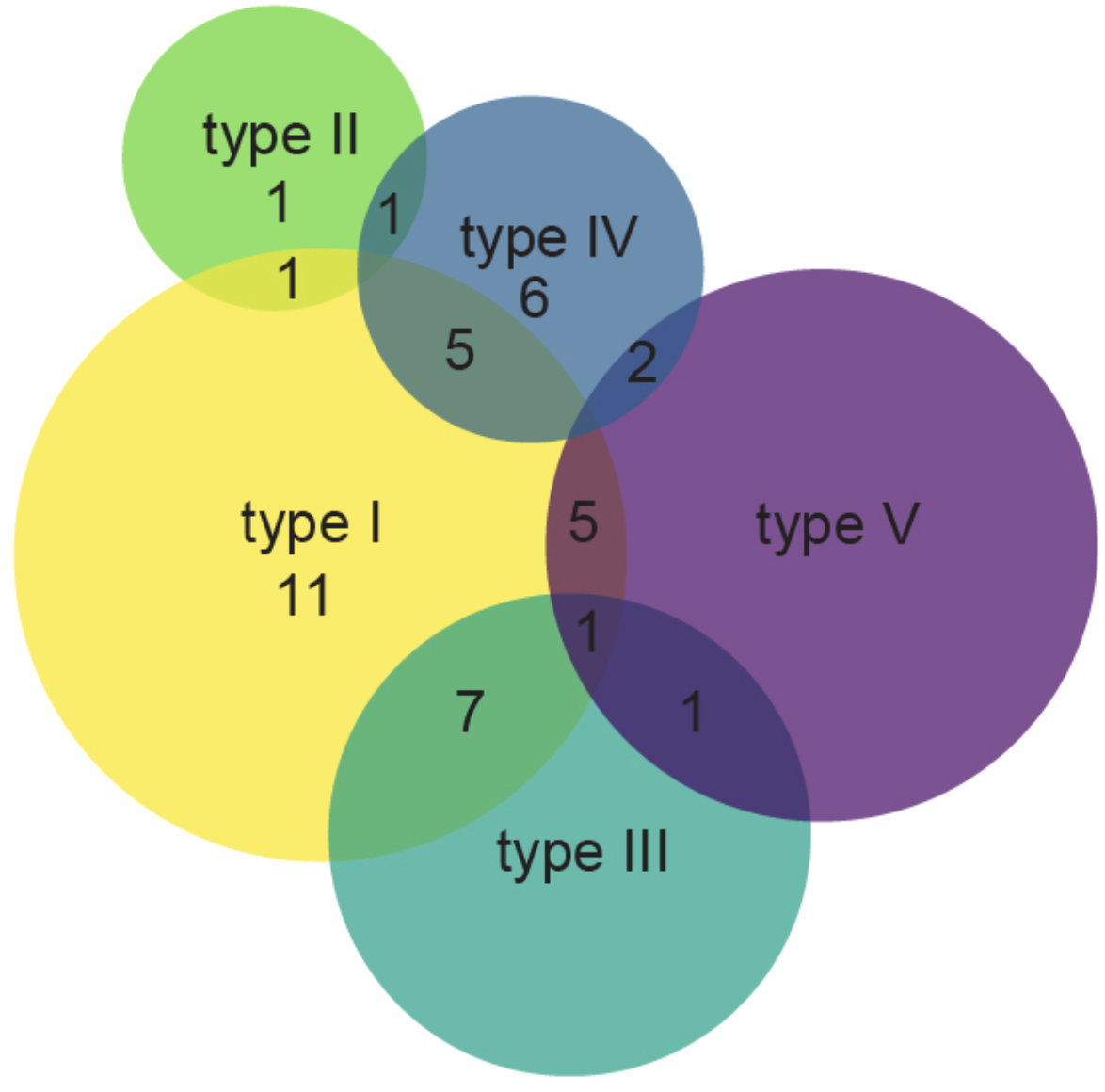
Co-occurrence of multiple CRISPR-Cas types on individual plasmids. The different CRISPR-Cas systems are color coded according to their type classification: I in yellow, II in light green, III in teal, IV in dark blue, and V in purple. The overlapping areas in the Venn diagram indicate co-occurrences of the different CRISPR-Cas types, where the number (n) of observations is indicated. For plasmids encoding type I systems, co-occurrence of different subtypes were also detected and the number of observations are indicated (n = 11).

**Supplementary Figure S6.**
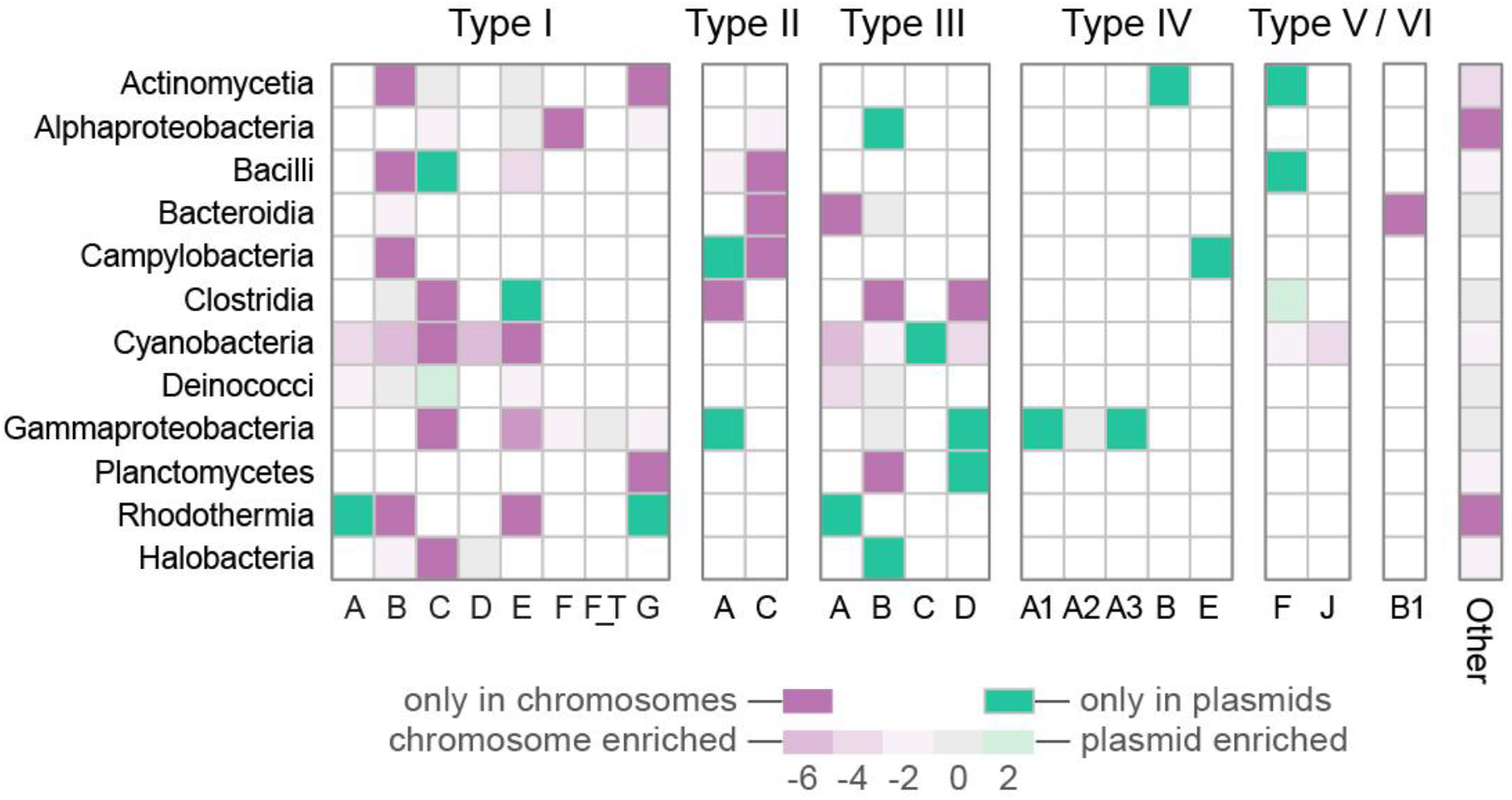
Direct comparison between CRISPR-Cas subtype prevalence across plasmids and their host-chromosomes (per taxa). Only plasmid-host chromosome pairs for which the plasmid and chromosome each contain at least one CRISPR-Cas are included in this analysis. Blank spaces represent subtypes for which no data is available. Color gradient denotes the log2 ratio between prevalence on plasmids and prevalence on chromosomes, such that positive values indicate plasmid enrichment and negative values indicate chromosome enrichment. The darkest shades of green and purple indicate subtypes only present in plasmids and chromosomes, respectively.

**Supplementary Figure S7.**
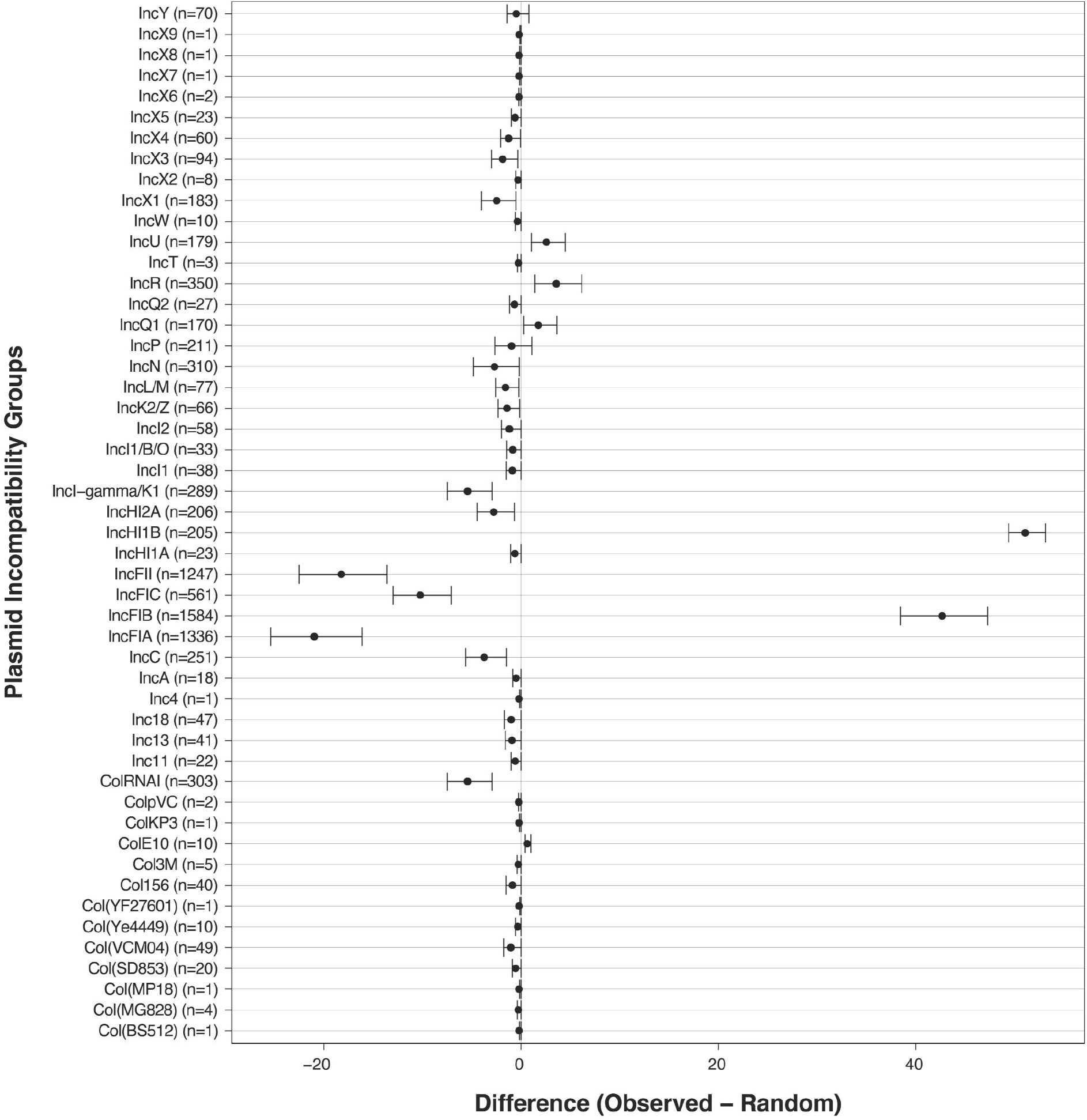
Distribution of CRISPR-Cas systems across plasmid Incompatibility (Inc) groups. Enrichment of CRISPR-Cas loci within plasmid incompatibility groups within the Inc-typeable fraction of the complete plasmid dataset. Single plasmids can belong to more than one Inc group. The observed distribution of CRISPR-Cas on Inc-typable plasmids was compared with a random distribution, in which CRISPR-Cas systems were placed in random Inc-typable plasmids. The mean and standard deviation from 1000 random permutations is shown. The difference is the number of observed CRISPR-Cas systems subtracted by the number of CRISPR-Cas systems in the random permutation.

**Supplementary Figure S8.**
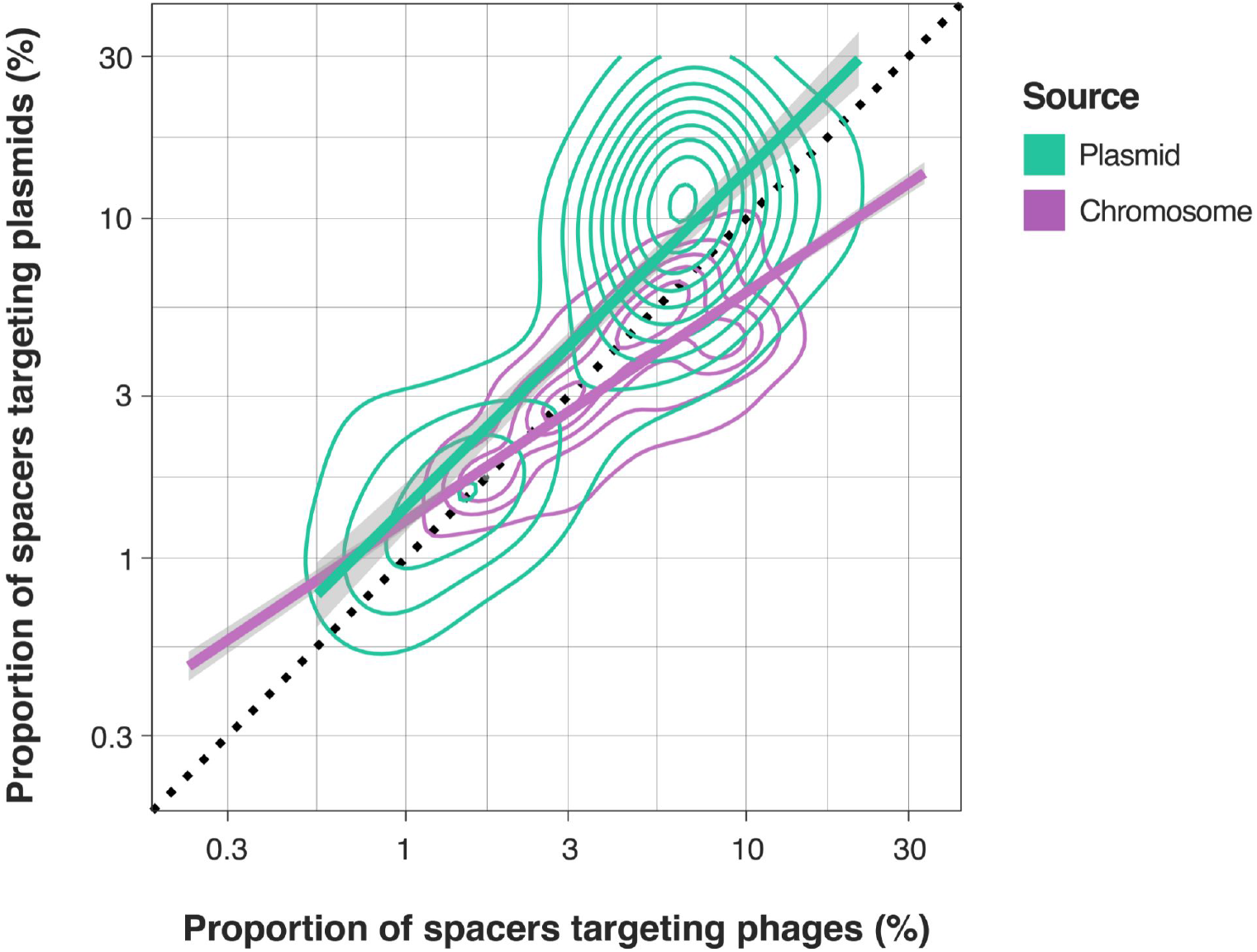
Proportion of spacers matching viral/plasmid sequences across all detected CRISPR arrays. Contour plot depicting the global targeting preference (plasmid vs. virus) for spacers within single arrays derived from plasmids (green) and plasmid-host chromosomes (purple). Regression lines with shaded areas correspond to 95% confidence intervals. The black dotted line indicates a 1:1 proportion of virus-vs plasmid-targeting spacers.

**Supplementary Figure S9.**
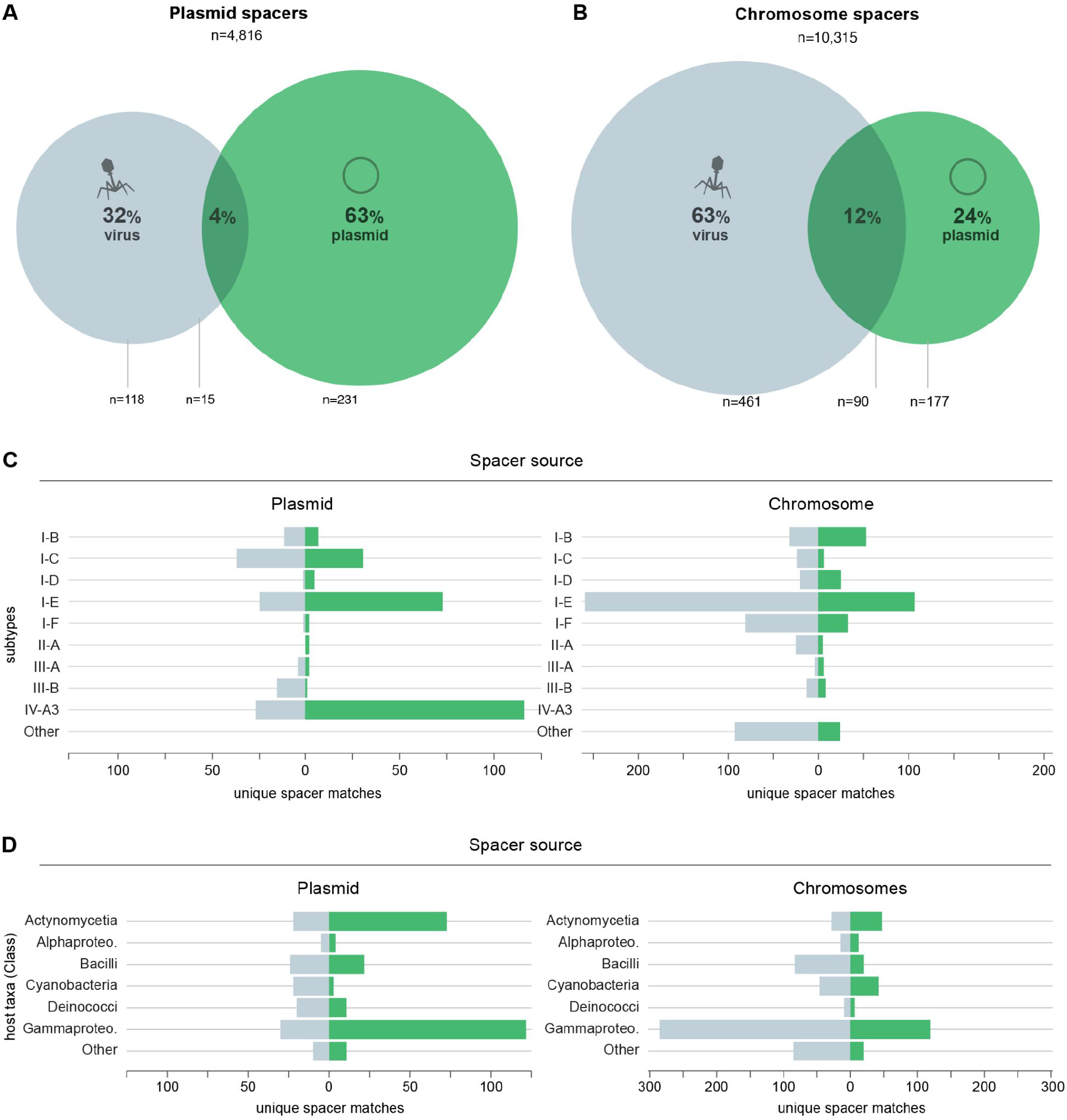
Direct comparison plasmid-host: only cells where both chromosome and (at least one) plasmid has CRISPR. The proportion of plasmid (**A**) and plasmid-host (**B**) chromosomal spacers matching plasmids (green) and viruses (grey). **(C)** Distribution of spacer-protospacer matches derived from plasmid (left) and plasmid-host chromosome (right) spacer contents, presented according to CRISPR-Cas subtype/variant and predicted spacer target: plasmids (green) and viruses (grey). Only the top 9 subtypes for both plasmids and chromosomes are represented, with the remaining grouped in “Other”. **(D)** Distribution of spacer-protospacer matches derived from plasmid (left) and host chromosome (right) spacer contents, broken down by host taxa and predicted protospacer origin: plasmids (green) and viruses (grey). Only the top 6 classes for both plasmids and chromosomes are represented, with the remaining grouped in “Other”.

**Supplementary Figure S10.**
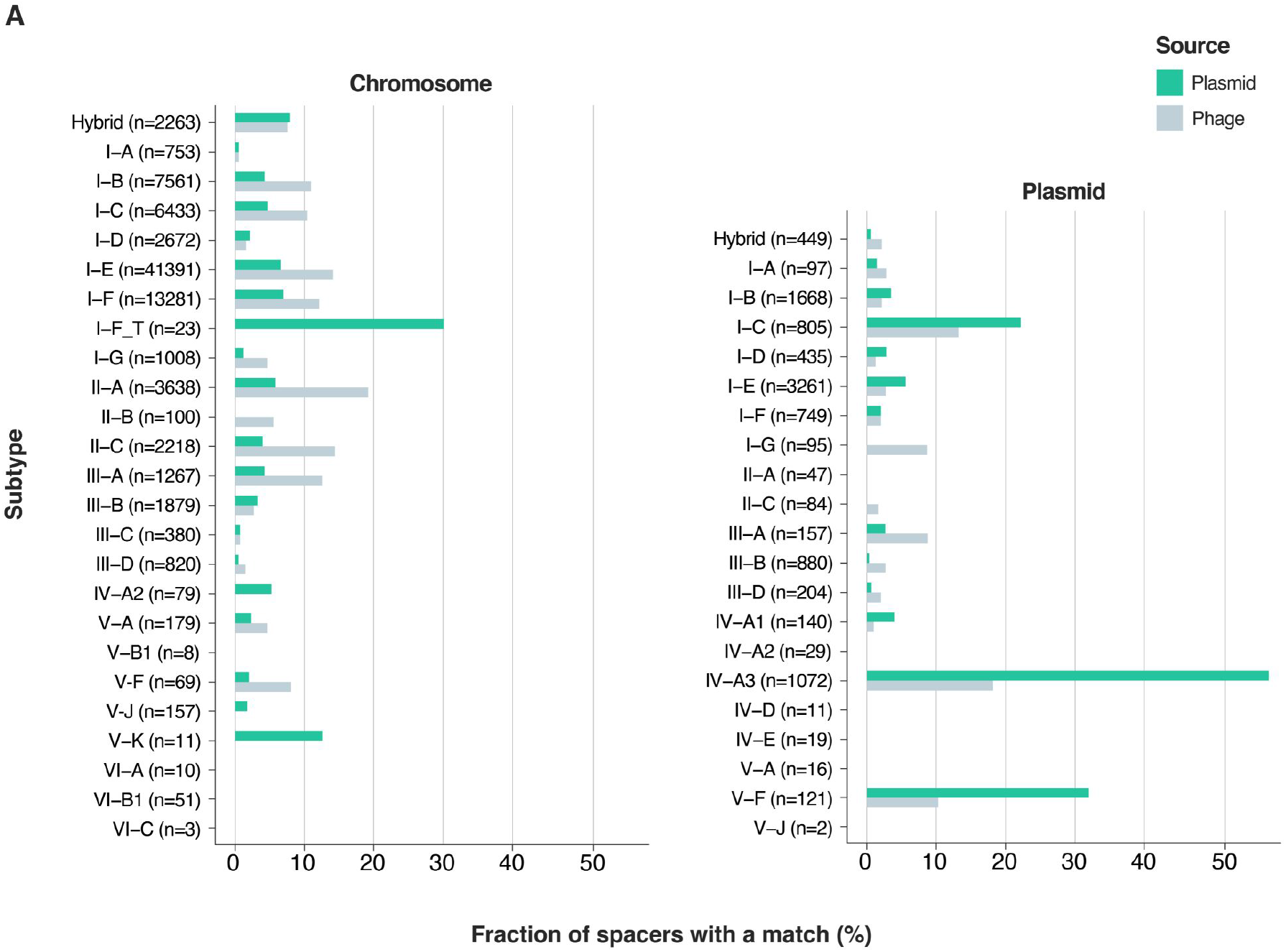
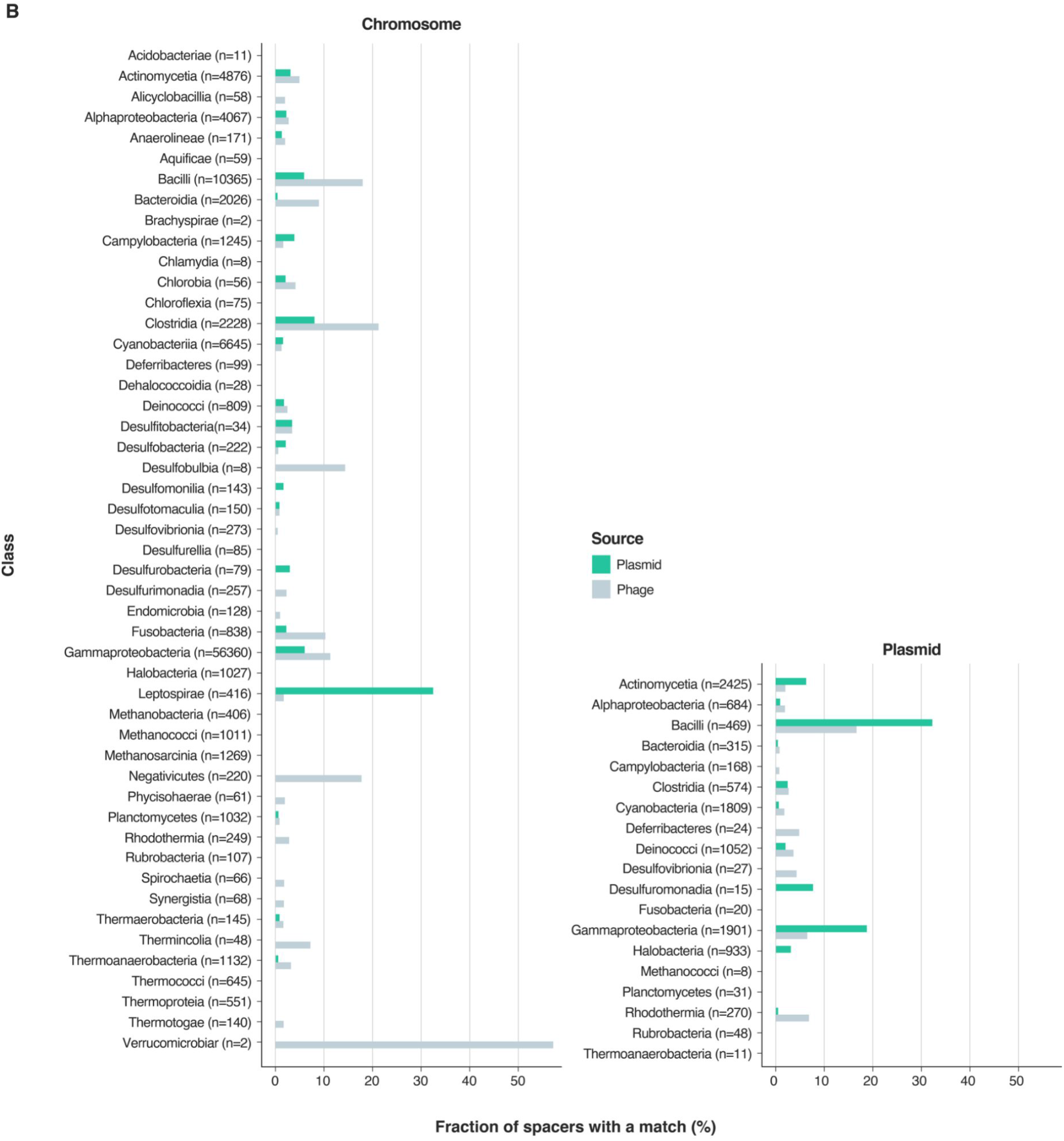
Fractions of unique bacterial spacer-protospacer matches per total number of spacers for each individual subtype and host class. **A)** Chromosomal and plasmid spacer-match distribution broken down by CRISPR-Cas subtype. **B)** Chromosome- and plasmid-derived spacer-match distribution broken down by host taxonomy (class level). Numbers (n=X) indicate the number of unique spacers.

**Supplementary Figure S11.**
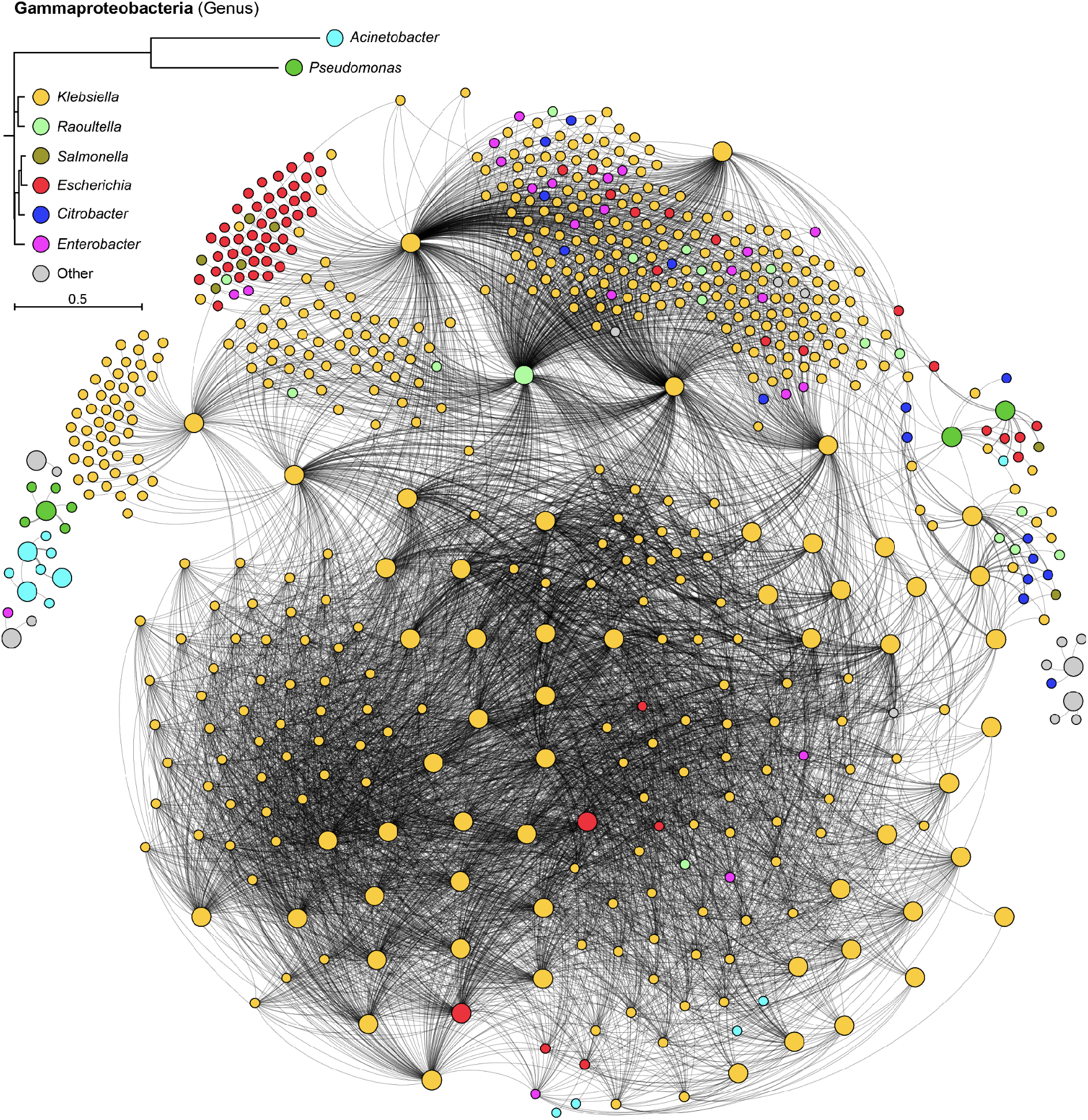
Clustering network of predicted plasmid-plasmid CRISPR-Cas targeting in Gammaproteobacteria. The plasmid-plasmid targeting network is colored at the host genus level, where nodes correspond to individual plasmids and edges represent predicted spacer-protospacer matches. The phylogeny in the legend is based on the median cophenetic distance from the GTDB whole-genome phylogeny, with the tree inferred by neighbor-joining. “Other” indicates plasmids without a known host, host with a different taxonomy than those displayed, or with a host with unspecific taxonomy. Large and small nodes indicate the presence or absence of CRISPR-Cas in the plasmid, respectively. Edge thickness is proportional to the number of spacer-protospacer matches between plasmid pairs.

**Supplementary Figure S12.**
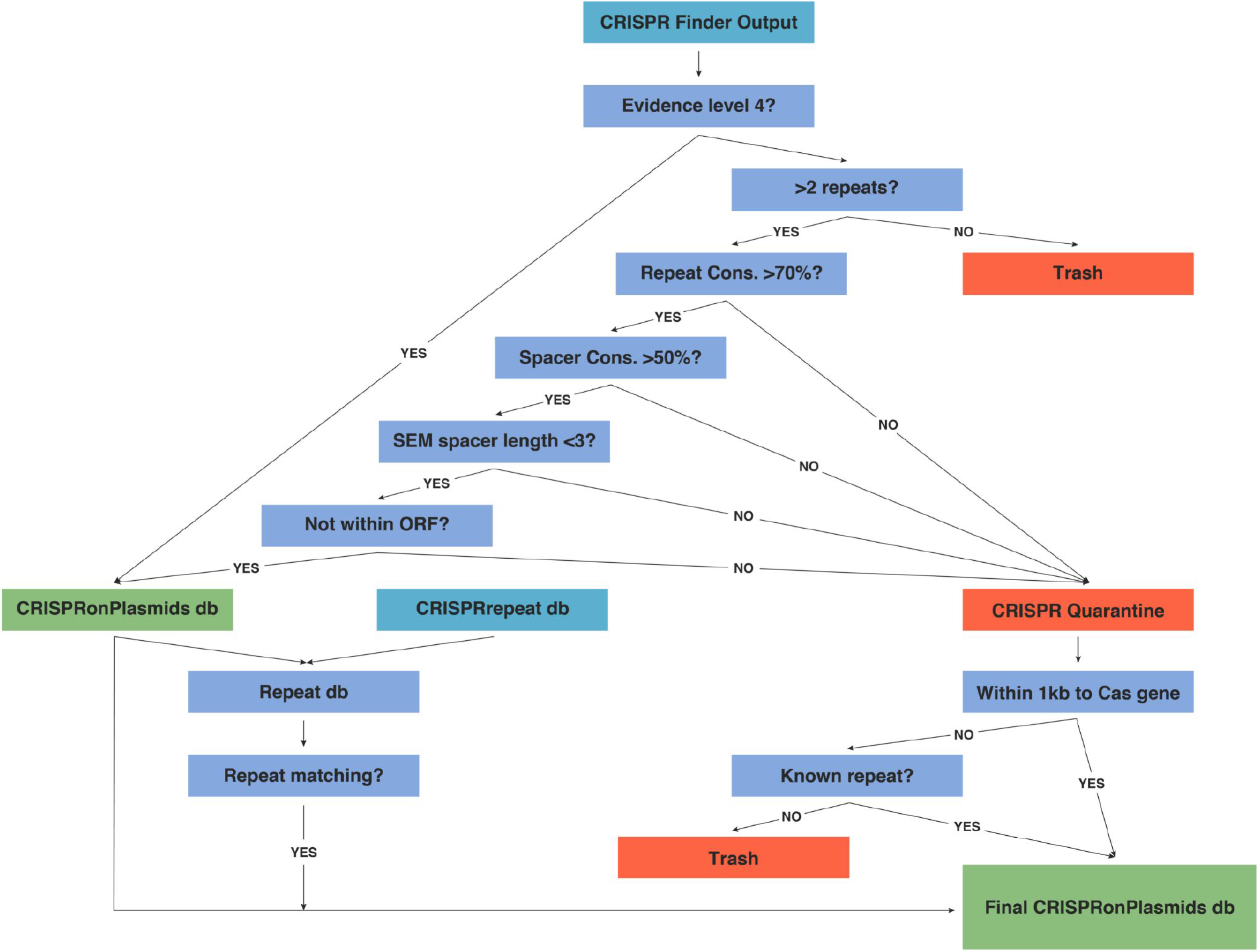
Decision tree used for elimination of false-positive arrays from the outputs of CRISPRCasFinder and inclusion of undetected CRISPRs. Briefly, high confidence arrays (evidence level 4) predicted by CRISPRCasFinder were automatically retained and included in the final CRISPRonPlasmids database (db). Arrays with lower evidence level scores were removed if the number of repeats was less than three; arrays with more than two repeats were placed into a quarantine list if the calculated average repeat conservation (cons.) across the array was higher than 70%, the spacer conservation was lower than 50%, the standard error of the mean (SEM) of the spacer lengths was less than 3, and if the array did not overlap with a predicted high confidence open reading frame (ORF). Putative arrays from the quarantine arrays were subsequently rescued for the analyses if they were located within 1 Kb of a predicted *cas* gene or if they matched (>95% coverage and identity) with high confidence CRISPR repeats (known repeats). Finally, the repeat db was blasted against plasmid/chromosome sequences to identify arrays that had been missed by the previous algorithm (repeat matching). Hits were considered true arrays when more than three repeats were identified (>95% identity and coverage), each within <100 bp from each other.

**Supplementary Figure S13.**
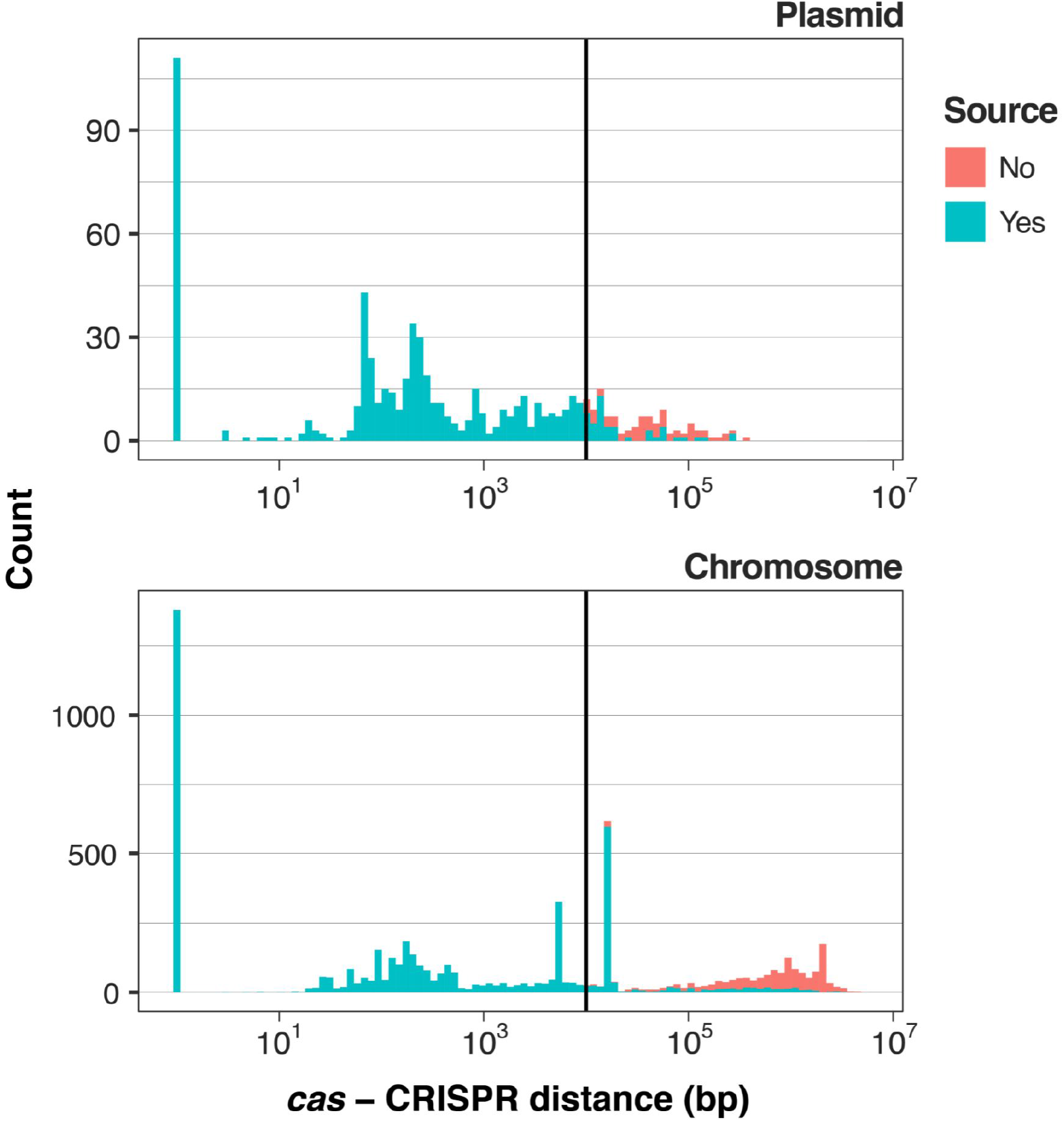
Distances between CRISPR arrays and the nearest *cas* operon. Only de-replicated plasmids and associated host chromosomes are included. All distances above 100 kb were grouped together in the 100 kb mark. Color denotes whether CRISPR arrays could be subtyped (blue; true) either by proximity to a *cas* operon (<10 kb threshold) or by repeat similarity with an array proximal to a *cas* operon (> 85% identity). Unassigned arrays are marked in red (false). Note that this does not include CRISPR arrays on contigs without any *cas* operon, which are the largest share of truly orphan arrays.

**Supplementary Figure S14.**
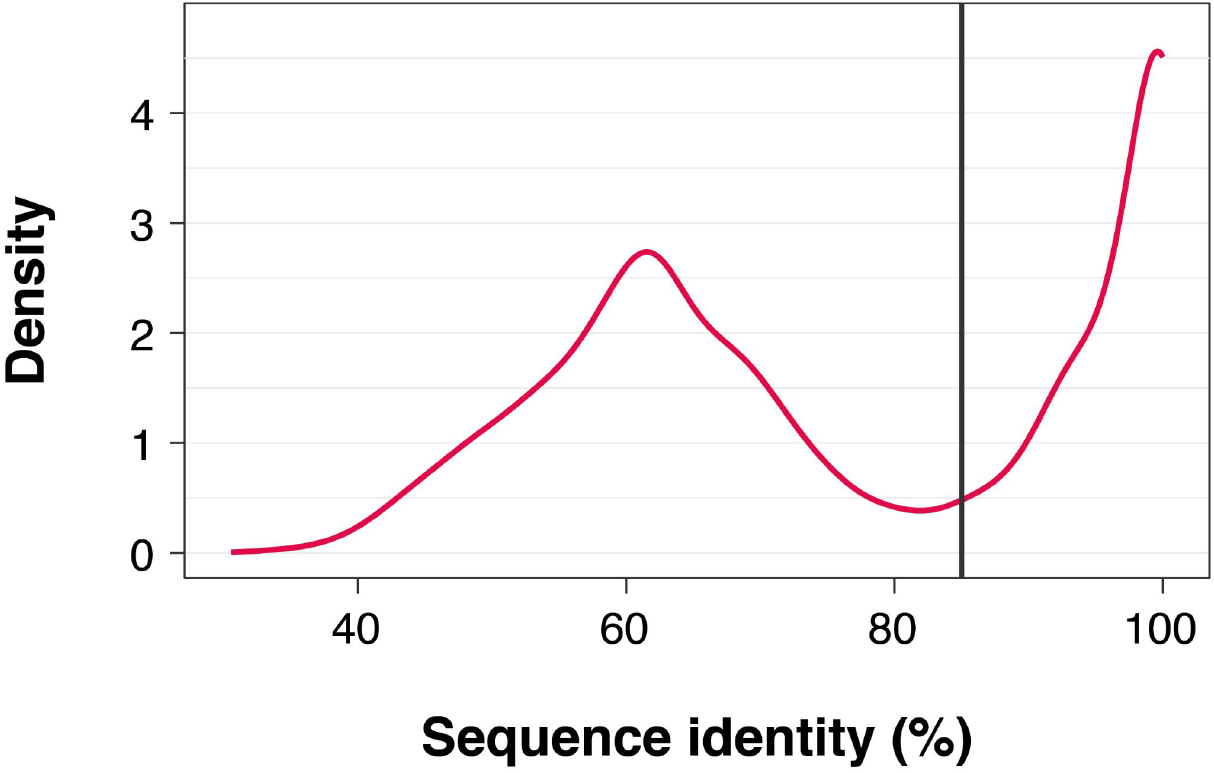
Density plot of sequence identities between consensus repeats of CRISPR arrays originating from the same contig. Vertical line shows the chosen cutoff for typing “distant CRISPR arrays”: orphan arrays that could be associated to a CRISPR-Cas locus within the same contig/genome.

## Notes

### Competing Interest Statement

The authors have declared no competing interest.

